# Cross-species convergence of surface polysaccharides shapes the virulence of an atypical carbapenem resistant uropathogenic *Escherichia coli* clone

**DOI:** 10.64898/2026.10.05.755231

**Authors:** Lachlan L. Walker, Zheng Jie Lian, Minh-Duy Phan, Nguyen Thi Khanh Nhu, Yvette Yu Ting Ong, Vitus Brix, Adam N. Marin, Brian S. Learman, Budi Permana, Irene Martinez-Roman, Samuel A.S. Morris, Kate M. Peters, Chelsea Stewart, Kay A Ramsay, David M.P. De Oliveira, Niels Pfennigwerth, Patrick N.A. Harris, David L. Paterson, Matthew J. Sweet, Leo A. Featherstone, Chelsie E. Armbruster, Brian M. Forde, Mark A. Schembri

**Affiliations:** Institute for Molecular Bioscience, The University of Queensland, Brisbane, Queensland, Australia; Australian Infectious Diseases Research Centre, The University of Queensland, Brisbane, Queensland, Australia; Department of Microbiology and Immunology, Jacobs School of Medicine and Biomedical Sciences, State University of New York at Buffalo, Buffalo, New York, USA; Frazer Institute, Faculty of Health, Medicine and Behavioural Sciences, The University of Queensland, Brisbane, Queensland, Australia; German National Reference Centre for Multidrug-resistant Gram-negative Bacteria, Department of Medical Microbiology, Ruhr-University Bochum, Universitätsstraße 150, 44801 Bochum, Germany; Pathology Queensland-Central Microbiology Laboratory, Queensland Health, Brisbane, Queensland, Australia; ADVANCE-ID, Saw Swee Hock School of Public Health, National University of Singapore, Singapore, Singapore; The Kirby Institute, Faculty of Medicine, University of New South Wales, Sydney, NSW, Australia; Evolution & Ecology Research Centre, School of Biological Earth and Environmental Science, UNSW-Sydney, Sydney, NSW 2052, Australia; School of Chemistry and Molecular Biosciences, The University of Queensland, Brisbane, Queensland, Australia

**Keywords:** uropathogenic *E. coli*, ST361, urinary tract infection, catheter, antibiotic resistance, AMR, capsule, virulence

## Abstract

Carbapenem-resistant *Escherichia coli* (CREC) is an emerging global health threat associated with limited treatment options and increased mortality. Here, analysis of 20,331 *E. coli* genomes from urinary tract infection (UTI) and urine samples identifies phylogroup A sequence type 361 (ST361) as an emerging CREC clone associated with UTI. Phylogenetic and phylodynamic analyses reveal two dominant ST361 clades that emerged in Asia around 2006 and subsequently disseminated intercontinentally. ST361 exhibits near pan-drug resistance, with *bla*_NDM-5_ conferring carbapenem resistance. In silico capsule and O-antigen typing reveals a highly conserved *Klebsiella*-like surface polysaccharide serotype in 95.9% of ST361 genomes, which we show contributes to virulence in UTI and catheter-associated UTI mouse models. Analysis of the mobile genetic elements harbouring *bla*_NDM-5_ in ST361 and other CREC further reveals a dynamic interplay between plasmid transfer and insertion elements that contributes to *bla*_NDM-5_ dissemination. Together, these findings identify ST361 as an emerging UTI-associated CREC clone and reveal the convergence of clonal expansion, virulence-associated surface architecture and mobile genetic elements in its spread.

## Introduction

Carbapenem-resistant *Enterobacterales*, including carbapenem-resistant *Escherichia coli* (CREC), represent a global health threat, designated as a critical priority pathogen by the World Health Organisation (WHO) due to their capacity to cause infections with limited treatment options, prolonged hospitalisation and increased mortality^1^. Carbapenem resistance in *E. coli* is commonly driven by enzymatic inactivation of the antibiotic, mediated by carbapenemases including the New Delhi metallo-β-lactamases (NDM), *Klebsiella pneumoniae* carbapenemases (KPC), oxacillinases (OXA), Verona integron-encoded metallo-β-lactamases (VIM) and imipenemases (IMP)^2^. Among these enzymes, NDM enzymes encoded by the *bla*_NDM_ gene are the most frequent cause of carbapenem resistance in CREC^3–5^. Directly relevant to carbapenem resistant Enterobacterales, mutations in penicillin-binding protein 3 (PBP3) represent an increasing threat, as these mutations can compromise the effectiveness of newer antibiotics developed specifically for their treatment, including cefiderocol and aztreonam-avibactam^6,7^.

CREC cause a broad spectrum of serious infections including urinary tract infections (UTI), bloodstream infections and device-associated infections^8^. Carbapenem-resistant infections primarily occur in healthcare settings^9,10^, with up to 80% of CREC infections attributed to healthcare exposure^10,11^. Among these infections, carbapenem resistance in uropathogenic *E. coli* (UPEC) is of particular concern, given that UPEC is the primary causative pathogen of UTI globally^12^. There are an estimated 404 million UTI cases annually^13^, with UPEC responsible for approximately 80% of these infections^12^. UTI can range from uncomplicated cystitis to life-threatening pyelonephritis and urosepsis^12^, with the acquisition of carbapenem resistance associated with increased treatment failure and high mortality^8^.

Global genomic surveillance of CREC has identified and characterised several sequence types (STs) associated with carbapenem resistance, including ST131, ST38, ST410, ST167 and ST405^4,5^. These STs span multiple phylogroups of *E. coli* with ST131 (B2), ST38 (D) and ST405 (D) belonging to phylogroups classically associated with extra-intestinal infection^14^ and characterised by high carriage of virulence factors^15–17^. In contrast, ST167 (A) and ST410 (C) are marked by reduced carriage of virulence factors but extensive antimicrobial resistance (AMR)^4,18^. Despite phylogroup A typically being associated with non-pathogenic intestinal colonisation^19^, CREC population studies identify phylogroup A as a major reservoir of carbapenem resistance genes^4^. Consistent with these findings, we recently identified a subclade of ST167 as an emerging atypical phylogroup A CREC lineage associated with UTI, characterised by resistance to carbapenems and third-generation cephalosporins, as well as reduced carriage of UPEC virulence factors, including the complete absence of type 1 fimbriae^18^. These findings raise the possibility that ST167 may not be an isolated exception but rather part of a broader emergence of phylogroup A lineages that contribute to the increasing burden of carbapenem-resistant UTI.

Here we analysed CREC from UTI and urine sources in EnteroBase^20^ and identified a previously under-characterised phylogroup A lineage, ST361, associated with human UTI. Phylogenetic analysis delineated two dominant carbapenem-resistant clades (clades IV and V) with a highly conserved *Klebsiella*-like KL25^Group^^1^:O9:H30 serotype. Bayesian phylodynamic and phylogeographic analyses indicated recent emergence around 2006, followed by intercontinental dissemination. Investigation of *bla*_NDM-5_ dissemination revealed a dynamic interplay between plasmid conjugation and transposable element-mediated structural variation. Wild-type strains from both clade IV and V efficiently colonised the urinary tract in a murine UTI model, with targeted mutation of the capsule genes resulting in reduced bladder colonisation. In contrast, a catheter-associated UTI (CAUTI) model revealed no role for the KL25^G^^1^ capsule in colonisation, but identified the O9 O-antigen as an important determinant of kidney colonisation. The contribution of O9 O-antigen to virulence was further supported by an *in vitro* whole-blood killing assay, which demonstrated that the O-antigen was essential for survival in human blood. Together, these findings identify ST361 as an emerging CREC lineage, with the presence of a *Klebsiella*-like capsule and O-antigen serotype contributing to the pathogenesis of this clone in niche-specific contexts.

## Results

### Major lineages of carbapenem-resistant *E. coli*

To identify the prominent lineages of CREC associated with UTI, we screened 20,331 genomes from human UTI and urine sources from EnteroBase^20^. *In silico* screening of antibiotic resistance genes found 3,427 *E. coli* genomes (16.9% of the dataset) harboured a carbapenem resistance gene; the *bla*_NDM-5_ gene was most common and present in 2,167 genomes (63.2%) (Fig. 1a). Phylogroup A comprised the largest proportion of CREC isolates (1,156 genomes), followed by phylogroup B2 (615 genomes) and phylogroup D (602 genomes) (Fig. 1b). In contrast, when we examined non-CREC isolates from UTI and urine sources we observed a dominance of phylogroup B2 (10,891 genomes) and D (2,321 genomes) (Fig. 1c). This latter distribution is more aligned with established UPEC literature that identifies strains from phylogroups B2 and D as the main cause of UTI^14^. Thus, within CREC from UTI-and urine-sources, there is a significant enrichment of phylogroup A *E. coli* isolates compared to non-CREC sourced from UTI-and urine-sources (Pearson’s χ² test = 2082.4, df = 1; *P*-value < 0.0001), with a markedly higher odds ratio of 7.09 (Fisher’s exact test; 95% confidence interval: 6.46-7.79).

**Figure 1.**
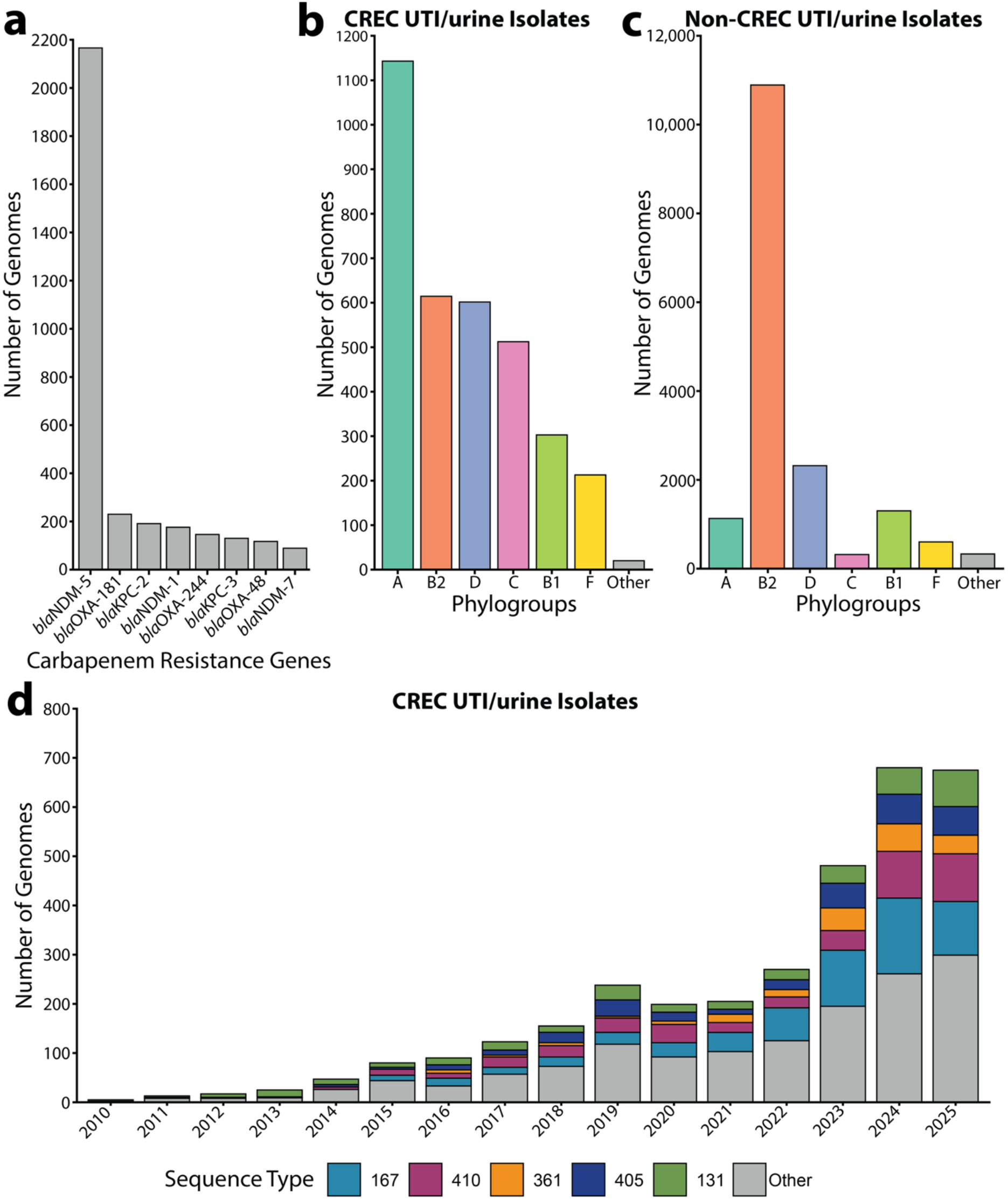
Overview of carbapenem resistance in human UTI/urine isolates from EnteroBase. (**a**) Bar chart of the number of genomes harbouring carbapenem resistance genes found in UTI and urine dataset from EnteroBase. Antibiotic resistance genes were identified with AMRfinderplus. The eight most frequently identified genes are shown. (**b**) Bar charts of carbapenem resistant genomes from UTI and urine sources in EnteroBase, colour-coded by phylogroup. (**c**) Bar charts of non-carbapenem-resistant genomes from UTI and urine sources in EnteroBase, colour-coded by phylogroup. “Other” in panels b and c includes phylogroup G and Clade I. (**d**) Stacked bar charts of carbapenem-resistant genomes colour-coded by sequence types over time (2010-2025). Coloured sequence types represent the five STs with the highest number of depositions in 2025. STs were determined using the MLST tool while phylogroups were determined using ClermonTyper.

**Supplementary Figure 1.**
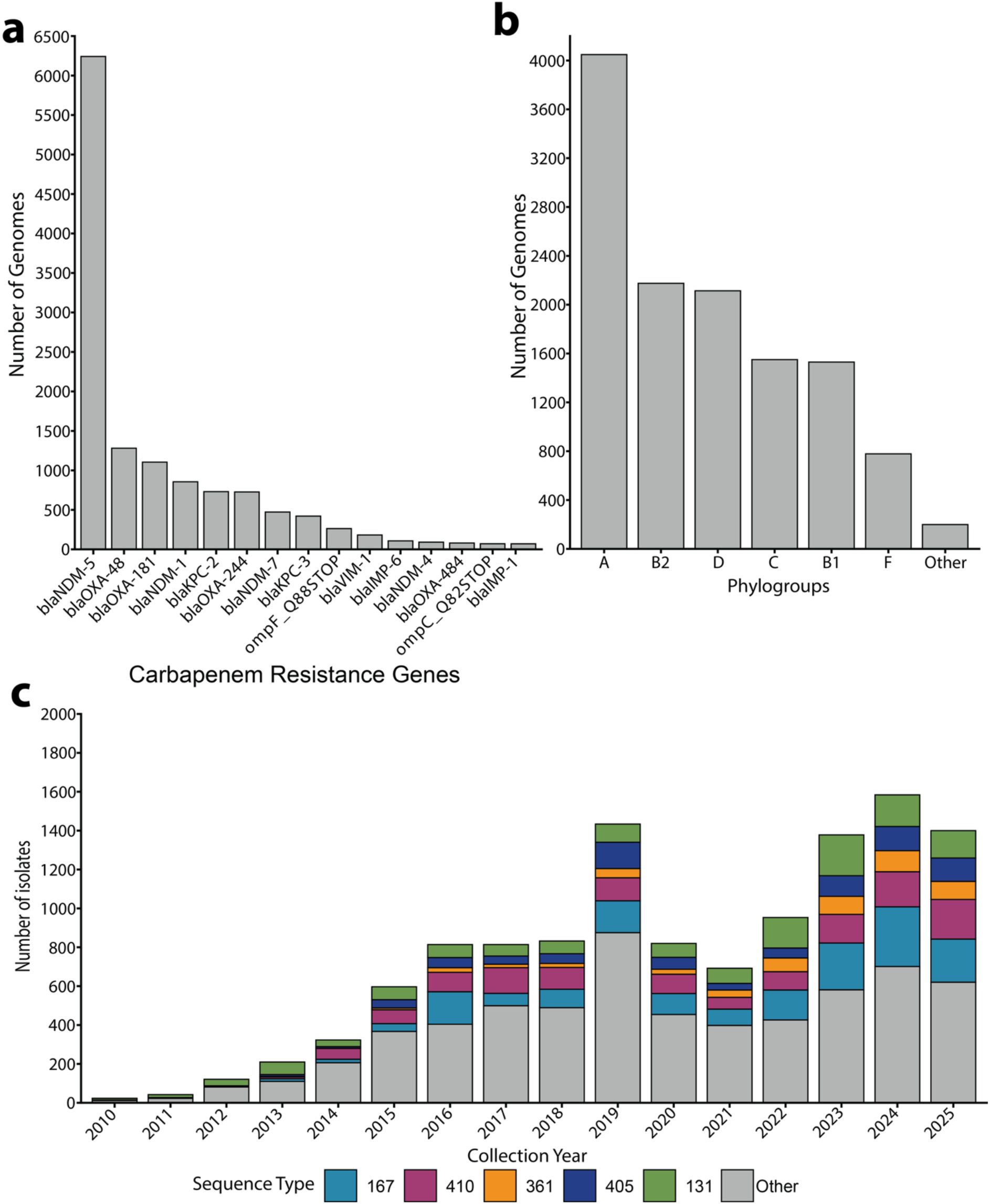
Overview of carbapenem resistance in human non-UTI/urine isolates from EnteroBase. (**a**) Bar chart of the number of genomes harbouring carbapenem resistance genes found in the dataset from EnteroBase. Antibiotic resistance genes were identified with AMRfinderplus (extracted from EnteroBase). The 15 most frequently identified genes are shown. (**b**) Bar charts of carbapenem-resistant genomes submitted to EnteroBase, organised by phylogroup. Other is representative of phylogroup G and Clade I. (**c**) Stacked bar charts of carbapenem-resistant sequence types over time (2010-2025). Coloured sequence types represent the five STs with the highest number of depositions in 2025. STs were determined using the MLST tool while phylogroups were determined using ClermonTyper.

**Supplementary Figure 2.**
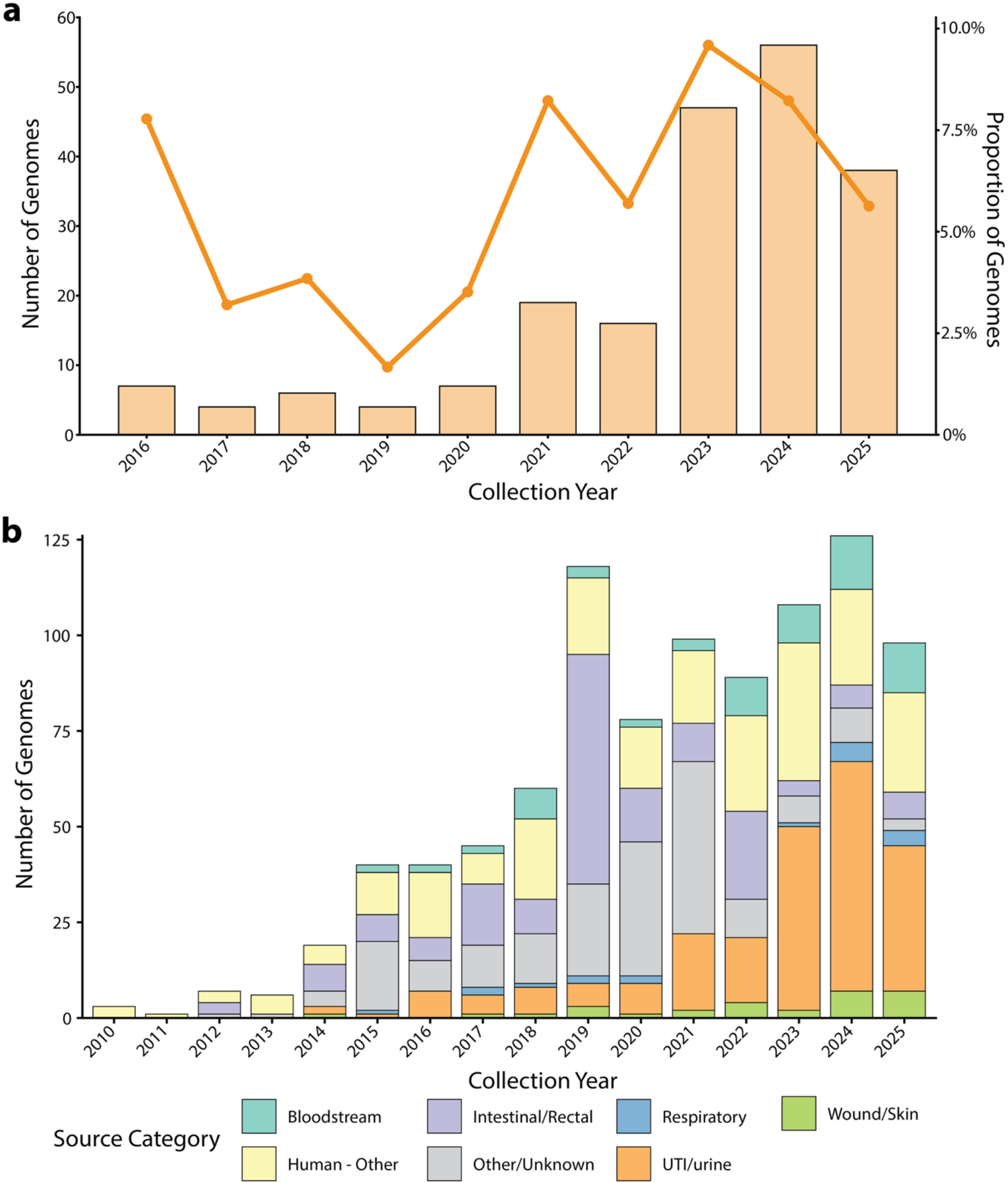
Isolation source metadata from EnteroBase for ST361. (**a**) Proportion of the UTI/urine dataset composed of ST361 genomes over time. The left y-axis shows the raw count of ST361 genomes identified in the UTI/urine dataset, plotted as a column graph by year. The right y-axis indicates the proportion of the dataset composed of ST361 isolates (line graph) as a percentage per year. (**b**) Bar chart of ST361 isolates deposited to EnteroBase over time; colours represent isolation source.

Analysis of depositions of CREC genomes to EnteroBase^20^ from 2010-2025 identified an increasing number of isolates over time (Fig. 1d). The STs with the highest number of genome depositions in 2025 included ST167, ST410, ST131, ST405 and ST361. Both ST167 and ST361 belong to phylogroup A, while ST410 (C), ST405 (D) and ST131 (B2) represent other phylogroups (Fig. 1d). These patterns were mirrored in CREC from non-UTI human sources, where the same five STs were deposited at the highest frequency (Supplementary Fig. 1).

The population structure and resistance determinants of several common *E. coli* clones that harbour carbapenem resistance genes have been described, including ST167^18,21–24^, ST410^25–28^, ST405^16,29^ and ST131^15,30,31^. However, comparatively little genomic information is available for ST361^5,32^. Between 2021 and 2025, the proportion of ST361 CREC genomes deposited per year showed that ST361 makes up 5-10% of CREC isolates from UTI and urine sources in EnteroBase^20^ (Supplementary Fig. 2). These findings are consistent with multiple independent single-centre, national and multi-national surveillance programmes spanning three continents, where ST361 was identified to account for 5-10% of CREC isolates in these studies (Supplementary Data 1)^16,21,32–39^. Together, these observations identify ST361 as a large yet under-characterised CREC lineage (Supplementary Data 1), prompting our detailed description of the population structure and genomic features of ST361.

### Phylogenetic analysis of ST361 reveals two dominant carbapenem-resistant clades

To investigate the population structure of ST361, we downloaded all publicly available ST361 genomes from EnteroBase^20^ (n = 981; Supplementary Data 2). A maximum-likelihood (ML) phylogenetic tree was constructed from 42,995 recombination-free core-genome single-nucleotide polymorphic (SNP) sites (Fig. 2). The phylogenetic structure of the ML tree was assessed by Bayesian analysis of population structure (fastbaps^40^). This analysis revealed five dominant clusters within ST361 that we defined as clades I to V that displayed high intra-cluster clonality and strong bootstrap support; the remaining clusters comprised 1-11 genomes (Supplementary Fig. 3a, Supplementary Fig. 3b). Clades IV and V were the largest, containing 380 and 415 genomes, respectively, with a median pairwise SNP distance of 67 core-genome SNPs (32 SNPs minimum; 130 SNPs maximum) (Supplementary Fig. 3c). Clade IV (175/380 genomes) and V (103/415 genomes) were significantly enriched for isolates derived from extra-intestinal sources (urine and blood) compared to clade I (2/95 genomes), clade II (3/36 genomes) and clade III (1/27 genomes) (Fisher’s exact tests with the Benjamini-Hochberg (BH) correction; Supplementary Data 3). Overall, these findings indicate that clades IV and V are disproportionately represented in the ST361 dataset, and the close phylogenetic relationship between them suggests a shared evolutionary background that may increase their pathogenic potential.

**Figure 2.**
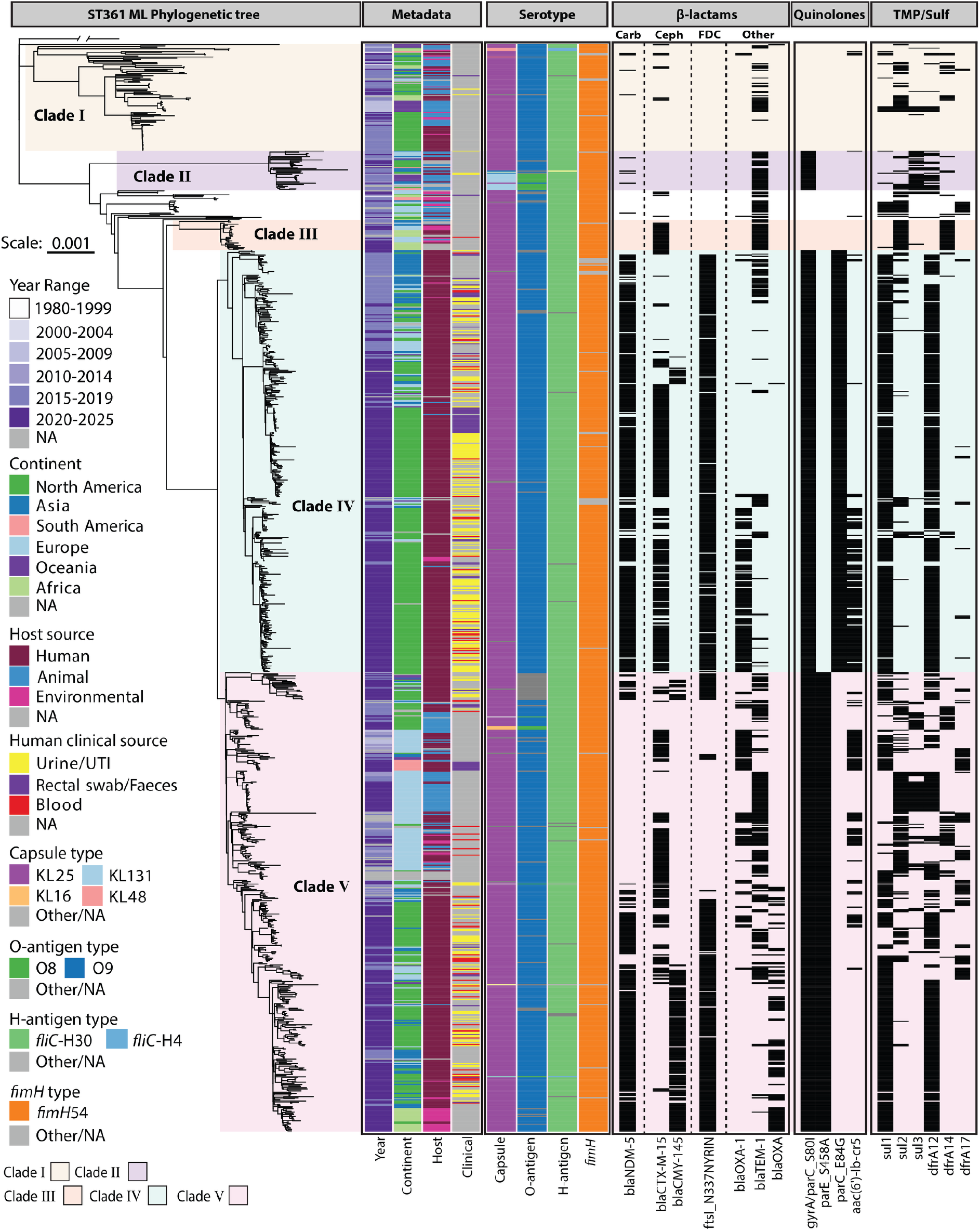
Maximum-likelihood phylogenetic tree of ST361. The phylogenetic tree was built from 42,995 recombination-free core-genome SNPs with IQ-TREE (GTR+G model and 1000 ultrafast bootstraps; ED1a (accession ID: CU928162) was used as the outgroup and MS25613 (accession ID: JCANXK000000000) was used as the reference genome). Fastbaps was utilised to determine the clade structure, and clades are shaded as per the key. Tree scale represents the number of nucleotide substitutions per site. Capsule type was determined with Kaptive 3.0 using the *Klebsiella* primary reference database. O-and H-antigens were identified using the EcOH database with ABRicate. The *fimH* antigen was determined by fimTyper. AMRfinderplus was used to identify antibiotic resistance genes. The annotation gyrA/parC_S80I represents the following mutations: gyrA_D87N, gyrA_S83L, parC_S80I. Carbapenem (Carb), cephalosporin (Ceph), cefiderocol (FDC) and trimethoprim/sulfonamide (TMP/Sulf) are abbreviated.

**Supplementary Figure 3.**
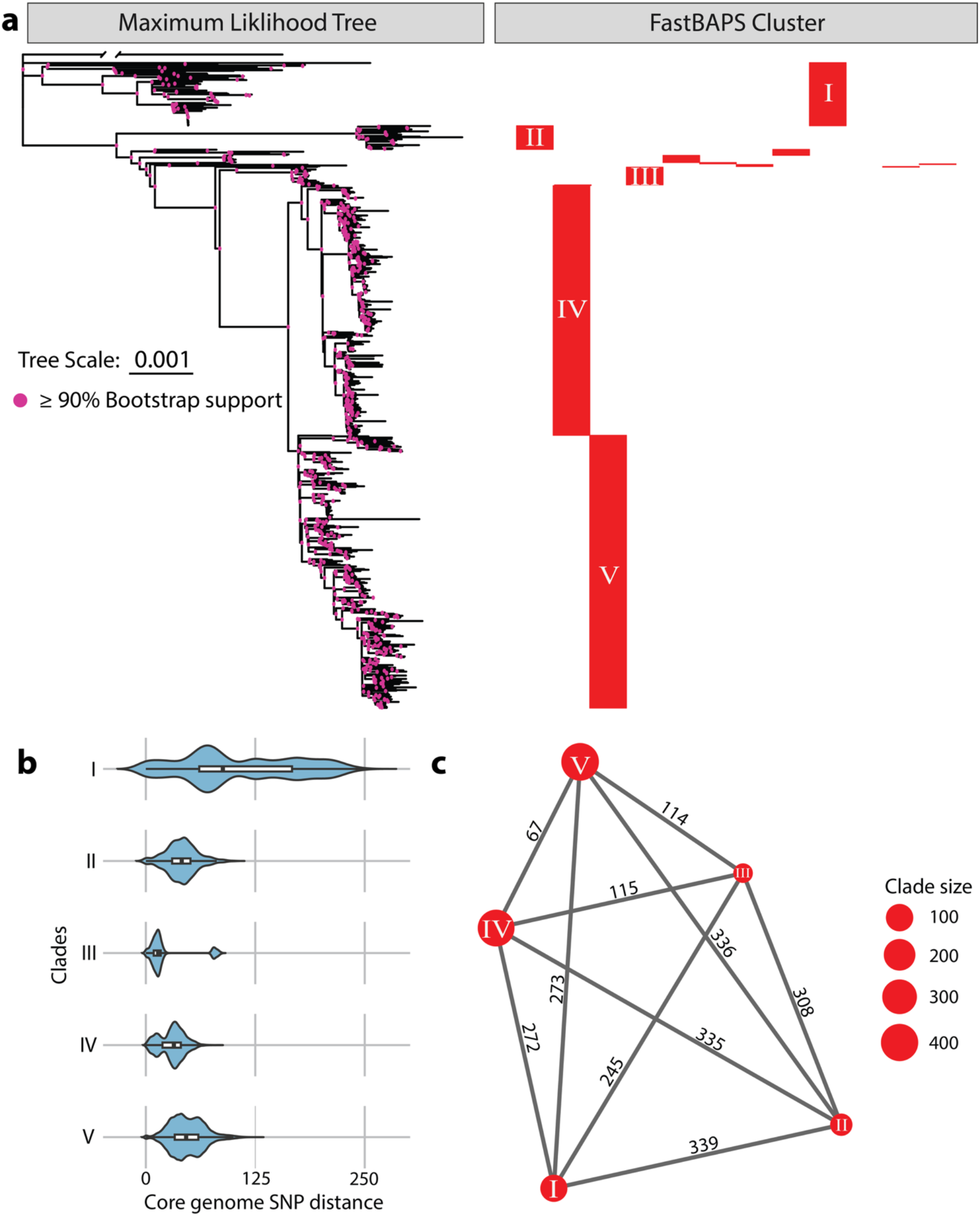
Maximum likelihood phylogenetic tree with Bayesian analysis of population structure and SNP distances between clades. (**a**) Nodes with more than or equal to 90% bootstrap support are indicated by pink dots. Designated clades are indicated in the panel on the right. (**b**) Violin plot visualising the pairwise core genome SNP distance of isolates within each clade. (**c**) Network plot visualising the median core genome SNP distance between isolates in each respective clade. The size of the dots indicates the number of genomes.

### Bayesian phylodynamic analysis reveals recent emergence of ST361 clades IV and V

We performed a Bayesian phylodynamic analysis with BEAST v1.10.4^41^ to reconstruct the demographic history of ST361 clades IV and V. This analysis used a Bayesian skyride coalescent tree prior^42^ to model the effective population size and an uncorrelated log-normal relaxed clock to model substitution rate (Supplementary Fig. 4a). We tested monophyletic constraint on both clades IV and V which produced comparable results, with the age of the root differing by approximately one year (Supplementary Fig. 4b); the clade V constraint was used for the primary model based on it containing the earliest sequenced isolates. Combined analysis of clade IV and V dated the most recent common ancestor (tMRCA) to 2006.4 (95% highest posterior density (HPD): 2005.7 – 2007.0) (Supplementary Fig. 4a). Individual analyses of clade IV and V suggest that clade V emerged first in 2006.4 (95% HPD: 2005.5 – 2007.2) followed by clade IV in 2011.3 (95% HPD: 2009.2 – 2012.9) (Supplementary Fig. 5). The clades exhibited similar evolutionary rates: posterior mean 7.26×10^-7^ substitutions/sites/year for clade IV (95% HPD: 6.16×10^-7^ – 8.41×10^-7^) and 6.89×10^-7^ substitutions/sites/year for clade V (95% HPD: 5.97×10^-7^ – 7.86×10^-7^).

**Supplementary Figure 4.**
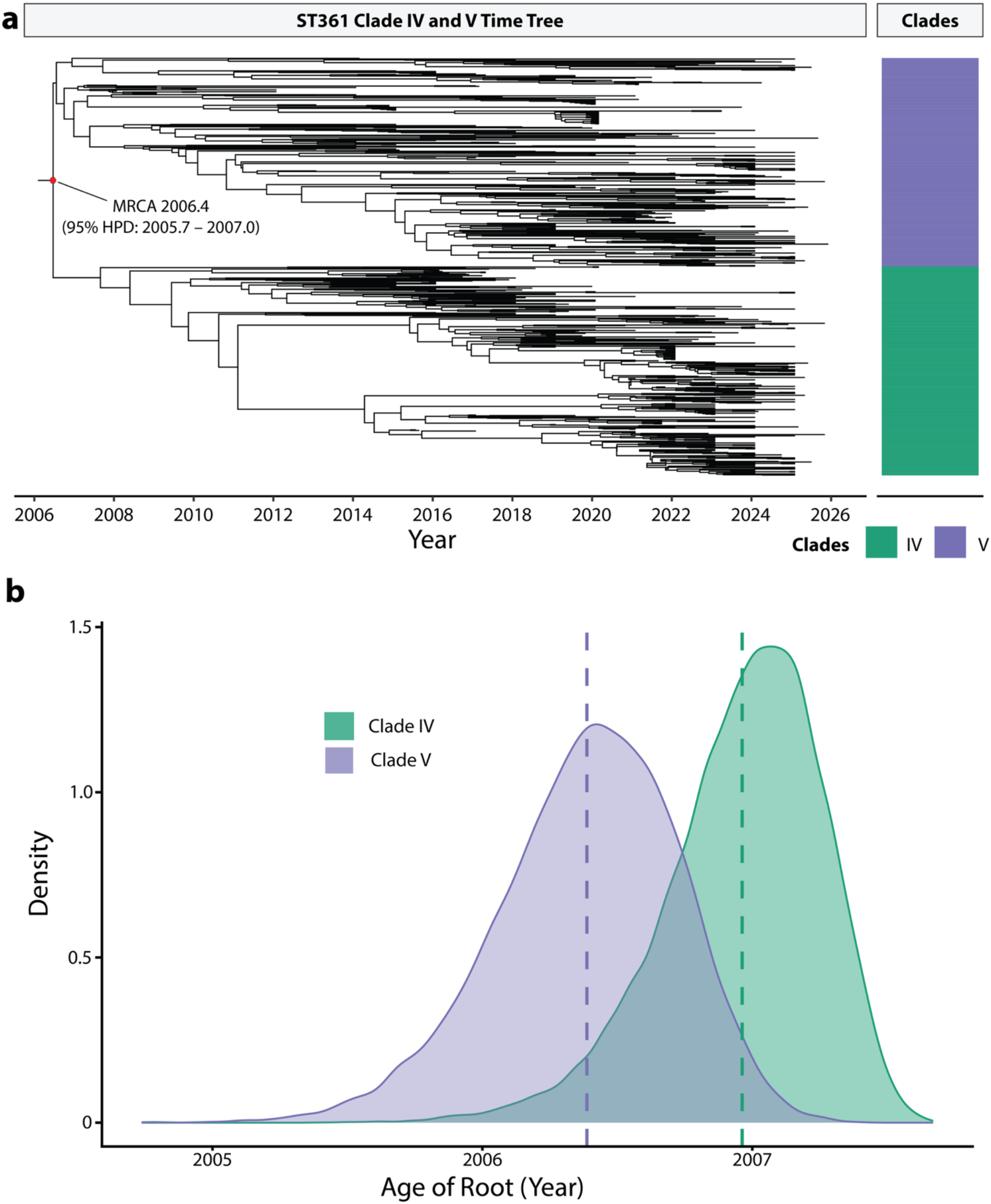
Bayesian analysis of ST361 clades IV and V. (**a**) A maximum clade credibility (MCC) tree of ST361 clades IV and V. Dataset consisted of 700 isolates, composed of 350 isolates from each clade, with MS25613 (accession ID: JCANXK000000000) used as reference. Red dot on the basal node of the tree represents the tMRCA of clades IV and V, annotated by the posterior mean tMRCA. (**b**) Density plot of the tMRCA of the combined clades IV and V analysis with differing monophyletic constraints applied. Green represents the analysis with clade IV under the monophyletic constraint and blue represents clade V. The dotted line depicts the posterior mean tMRCA for each clade.

**Supplementary Figure 5.**
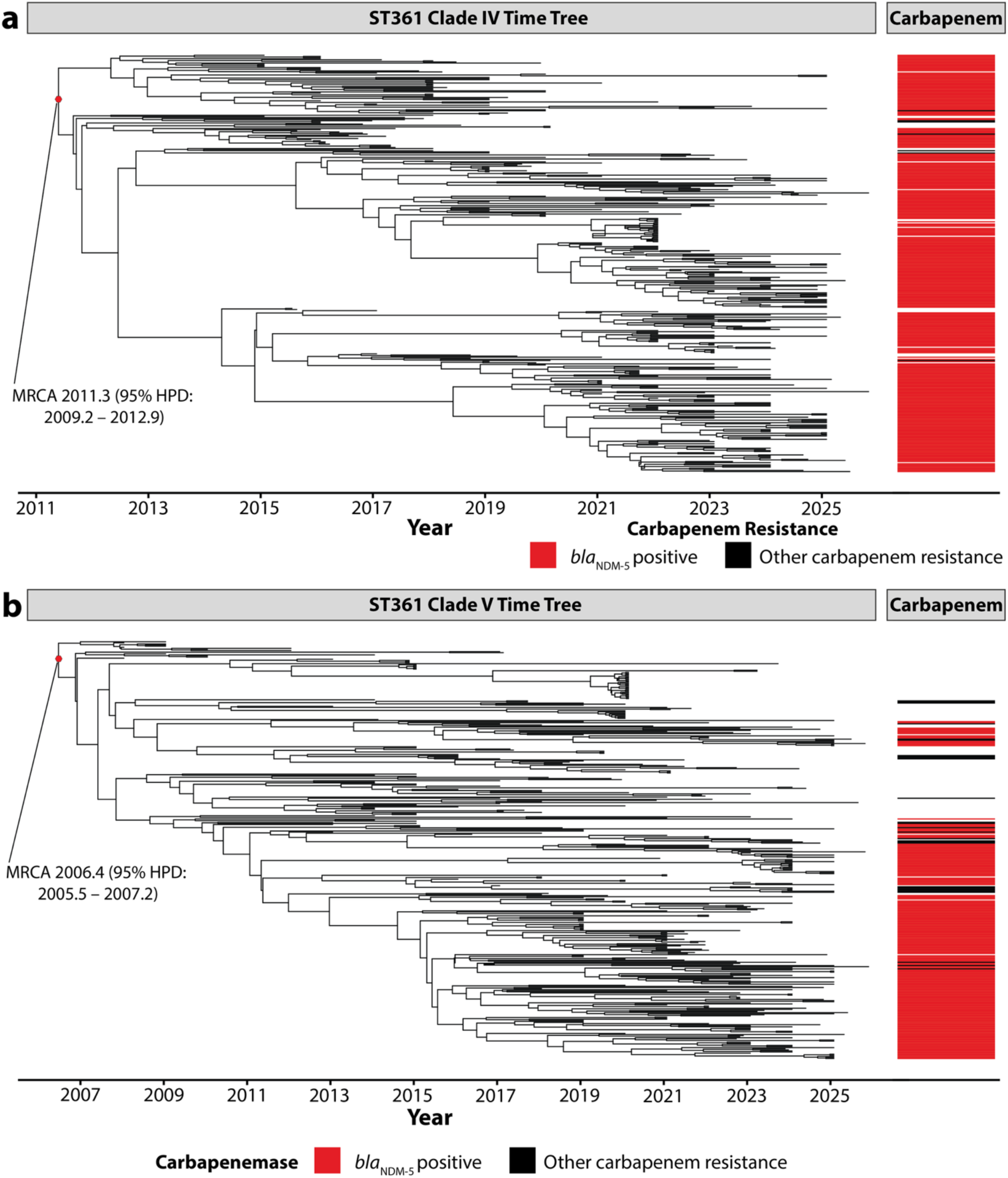
Bayesian MCC phylogenetic tree of ST361 clades IV and V. Time-scaled MCC trees of ST361 clade IV. (**a**) and V (**b**) containing 376 and 382 genomes respectively. Red dot indicates the predicted tMRCA. The heatmap on the right depicts genomes positive for *bla*_NDM-5_ and other carbapenem resistance genes.

### Distinct population trajectories and intercontinental spread for clades IV and V

Bayesian skyride analysis revealed similar but temporally separated trajectories for effective population sizes of clade IV and V. Between 2007-2012, the increasing effective population size (a measure of diversity) of the combined analysis reflected expansion of clade V prior to the emergence of clade IV (Fig. 3a). Clade V exhibited a modest expansion from 2015-2020, after which the effective population size stabilised and remained relatively constant through to 2025 (Fig. 3a). Conversely, clade IV expanded from 2011-2017 and showed a minor contraction in effective population size during the COVID-19 pandemic (2020-2022), before a subsequent recovery and growth until 2025 (Fig. 3a).

**Figure 3.**
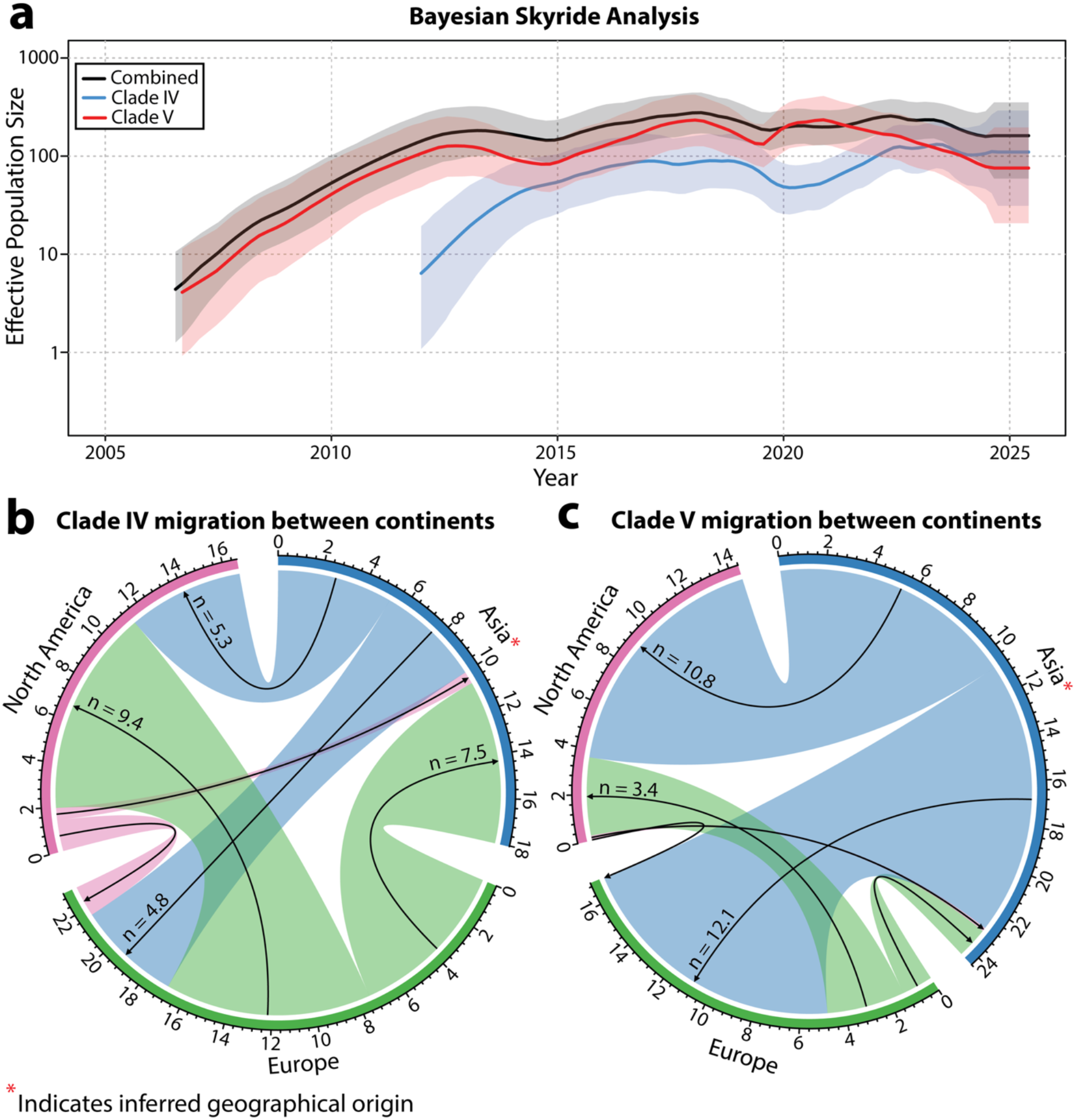
Bayesian skyride plots and phylogeographic transitions of ST361 clades IV and V. (**a**) Bayesian skyride graph depicting changes in effective population size over time. The three analyses are presented with the colour corresponding to the distinct clade IV, V and combined analyses. Shading indicates the 95% highest posterior density for effective population size on a log scale. (**b**, **c**) A chord diagram indicating the posterior mean number of transitions pre-2022 for clade IV (**b**) and V (**c**). Arrow direction indicates source-to-sink dynamics; numbers indicate posterior mean number of transition events between regions. Red asterisk indicates inferred geographical origin.

We next employed a Bayesian phylogeographic model to infer the geographic origin and dispersal patterns of clades IV and V before 2022 to mitigate geographical sampling bias in recent years. Ancestral state reconstruction inferred that both clades IV and V emerged in Asia with posterior proportion of 0.74 and 0.90, respectively. Analysis of Markov jumps for clade IV identified frequent transition events from Europe to Asia (mean 7.5; 95% HPD: 2 – 20) and Europe to North America (mean 9.4; 95% HPD: 5 – 16) (Fig. 3b). Support for other transition events was more diffuse, with transition events from Asia to Europe (mean 4.8; 95% HPD: 0 – 8) and Asia to North America (mean 5.3; 95% HPD: 0 – 10) (Fig. 3b). In contrast, clade V showed a pattern of asymmetrical spread originating from Asia with common transition events from Asia to Europe (mean 12.1; 95% HPD: 6 – 16) and Asia to North America (mean 10.8; 95% HPD: 7 – 13) (Fig. 3c). Overall, these phylogeographic analyses suggest that despite the high representation of North American isolates in recent years, both clades IV and V were likely introduced to North America through multiple transition events, underscoring their capacity for intercontinental spread.

### ST361 exhibits high AMR gene carriage consistent with a multidrug resistance phenotype

Comparative analysis of *in silico* AMR genes of ST361 isolates revealed clades IV and V harbour a significantly higher mean number of AMR genes per genome (18.5 and 19.6, respectively; Supplementary Data 4), compared to clades I (7.8), II (16.8) and III (12.3) (Pairwise Wilcoxon rank-sum tests with BH correction, *P* < 0.05; Supplementary Data 5). Distinct clade specific patterns of AMR gene carriage were also observed (Supplementary Fig. 6). For example, differences in quinolone resistance-associated point mutations were evident between clades IV and V. Clade IV carried the ParC*_*E84G mutation, whereas clade V harboured the ParE_S458A mutation (Fig. 2). Clades IV and V also exhibited high carriage of resistance genes against carbapenems (90.2% and 51.1% of genomes, respectively; Fig. 2). The most frequently detected carbapenem resistance gene in ST361 was *bla*_NDM-5_, present in 57.5% (564/981) of genomes. Treatment options for carbapenem-resistant *Enterobacterales* include cefiderocol and aztreonam-avibactam. Concerningly, we identified that 94.9% (535/564) of ST361 isolates harbouring *bla*_NDM-5_ also contained the cefiderocol and aztreonam-avibactam resistance-associated YRIN insertion in the PBP3 binding protein (FtsI_N337NYRIN)^6,7^.

To test if the co-occurrence between the FtsI_N337NYRIN mutation and the *bla*_NDM-5_ gene conferred phenotypic non-susceptibility to cefiderocol, we performed MIC analysis on nine clade IV and two clade V ST361 isolates carrying both determinants that were available in our collection. In total, 7/11 isolates (6/9 clade IV and 1/2 clade V) were non-susceptible with MIC > 2 mg/L (Supplementary Data 6). Although our analysis was limited by sample size, these findings are supported by previous reports of cefiderocol non-susceptibility in ST361 isolates^32,43^. While we did not directly assess the susceptibility of ST361 isolates to aztreonam-avibactam in this study, other studies have reported that ST361 isolates are non-susceptible to this treatment option, likely owing to both the YRIN insertion in the PBP3 binding protein and production of AmpC β-lactamases^7,44,45^. We also assessed the broader phenotypic susceptibility of these isolates across eight antibiotic classes (Supplementary Data 7). The majority of isolates (9/11) from clades IV and V were non-susceptible to all antibiotic classes tested except for aminoglycosides and nitrofurantoin. For aminoglycosides, approximately half of the isolates remained susceptible to amikacin, gentamicin and tobramycin (Supplementary Data 7). Overall, these findings indicate a broadly multidrug-resistant phenotype, with limited treatment options available, including with recently developed antibiotics.

**Supplementary Figure 6.**
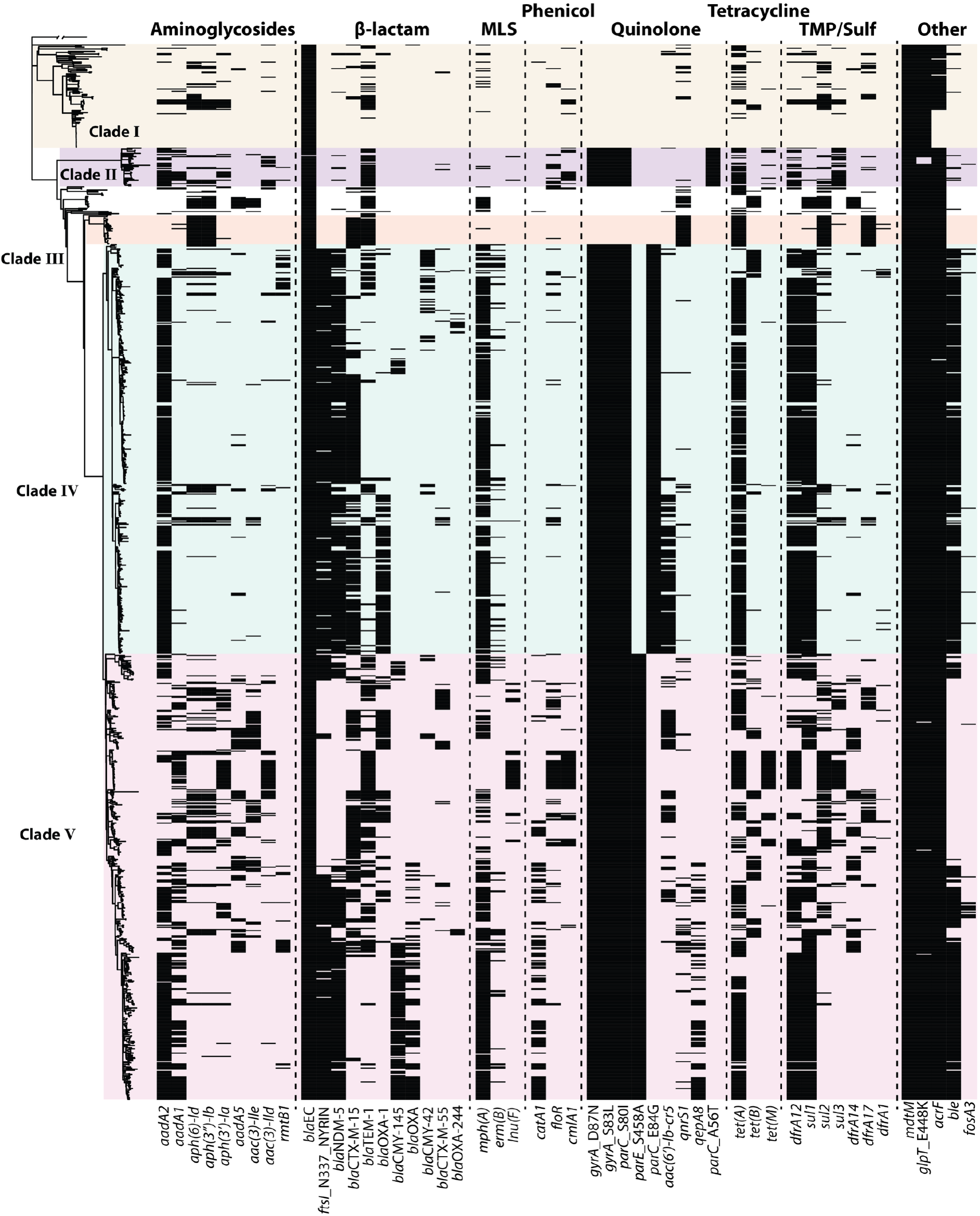
Presence and absence plot of AMR genes harboured in ST361. Maximum-likelihood phylogenetic tree of ST361 with a heatmap of the 50 most frequently identified antibiotic resistance genes. Antibiotic resistance gene carriage (black bars) were identified with AMRfinderplus, coloured shading across the phylogeny indicates the clades defined by Fastbaps. Macrolide-lincosamide-streptogramin (MLS); Trimethoprim-sulfamethoxazole (TMP/Sulf).

### Acquisition and spread of *bla*_NDM-5_ in major carbapenem-resistant lineages

The high proportion of isolates in clade IV and V harbouring *bla*_NDM-5_ prompted us to investigate the acquisition of this resistance determinant in ST361 (Fig. 2; Supplementary Fig. 5). To explore this, we screened 5,052 complete *E. coli* genomes (from RefSeq; accessed: 22/05/2026) which included 55 ST361 genomes (11 clade IV and V isolates sequenced in this study and 44 publicly available). In total, we identified 443 *bla*_NDM-5_ sequences from 414 complete *E. coli* genomes (of which 42 *bla*_NDM-5_ sequences were identified in 38 ST361 genomes), with two primary structural variants in the 10kb region immediately downstream of each *bla*_NDM-5_ copy (Fig. 4a; Fig. 4b), determined using a splitting tree as previously described^46^.

**Figure 4.**
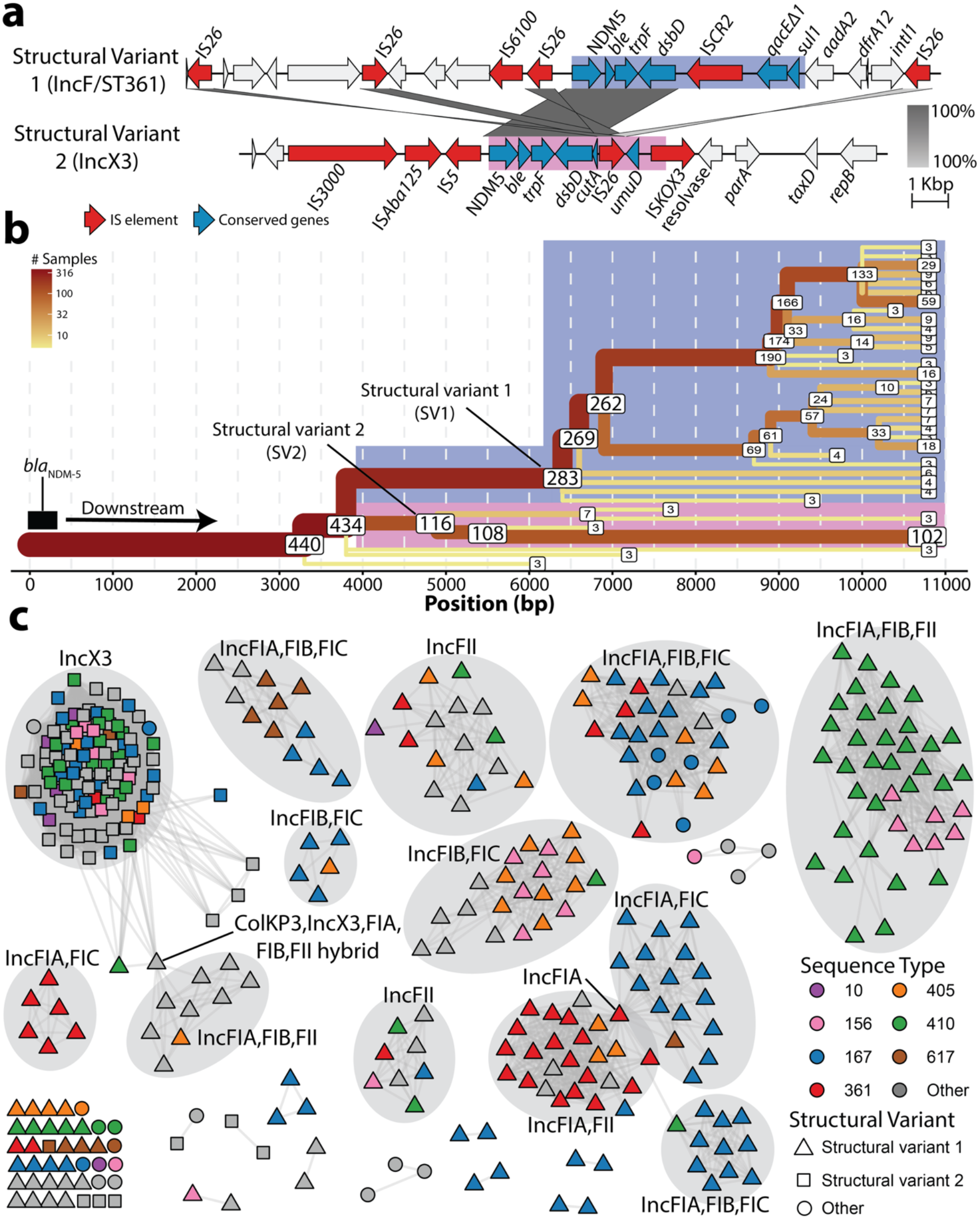
Structural conservation and plasmid diversity drive *bla*_NDM-5_ spread across high-risk *E. coli* lineages. (**a**) Easyfig plot depicting structural variant one and two. The CDS denoted in red represent different IS elements, the blue CDS represent genes that are part of the conserved module for each respective structural variant. MS25613 was used as a representative sequence for structural variant one and NZ_CP034965.1 was the representative sequence for structural variant two. Shading in blue and pink defines the “core” region identified in the respective structural variants. The x-axis represents distance from *bla*_NDM-5_. (**b**) A splitting tree of the structural variants identified downstream of *bla*_NDM-5_ from the *E. coli* RefSeq database. The labels on the node represent the remaining number of contigs on each branch, with branching points containing less than three contigs not shown. Shading in blue and pink represent the structural variants 1 and 2, respectively, and derivatives of the structural variants. (**c**) Plasmid clustering of *bla*_NDM-5_ positive plasmids from the *E. coli* RefSeq database with pling. Only the largest community, containing 375/402 of the *bla*_NDM-5_ plasmids, is visualised. Nodes represent plasmids, with edges indicating a double cut and join score ≤ 4 within each community as determined by pling. Colour of node indicate the ST of specific *E. coli* lineages. Grey shading of certain subcommunities denotes the major plasmid replicon types harboured, as identified by the plasmid finder database.

**Supplementary Figure 7.**
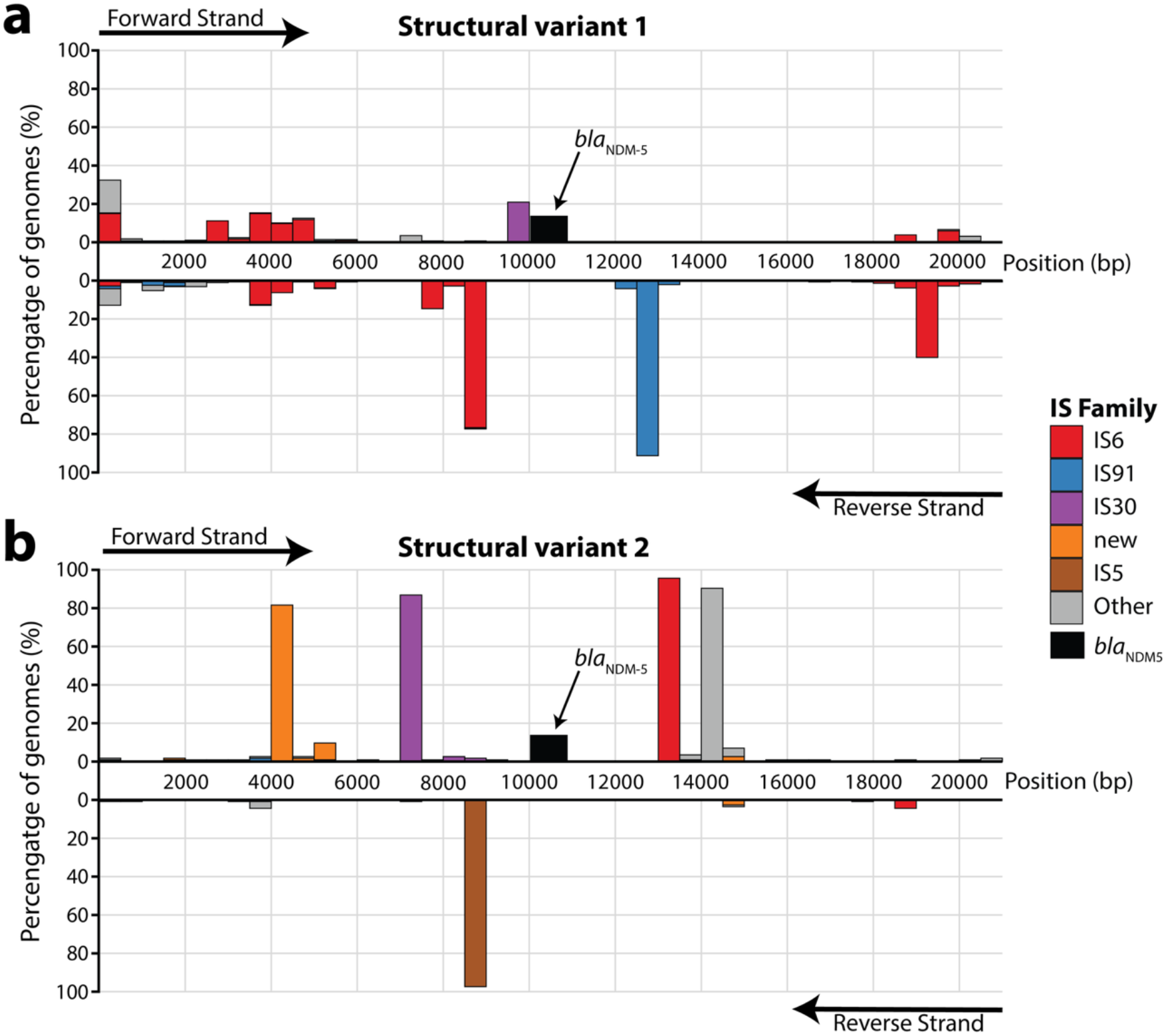
Analysis of *bla*_NDM-5_ genomic loci. (**a,b**) Distribution of IS elements identified 10 kb up and downstream of the *bla*_NDM-5_ gene from the *E. coli* RefSeq database in structural variant 1 (a) and structural variant 2 (b). The x-axis represents position (bp) with *bla*_NDM-5_ beginning at 10 kb. Plotted columns represent the start position of IS elements in 500 bp bins. Bars above the x-axis are IS elements identified on the positive strand, bars below the x-axis are IS elements identified on the negative strand. Arrows indicate orientation of the strand. Bars are coloured by IS family. The position of *bla*_NDM-5_ is indicated by a black box. IS elements were identified with ISEscan.

**Supplementary Figure 8.**
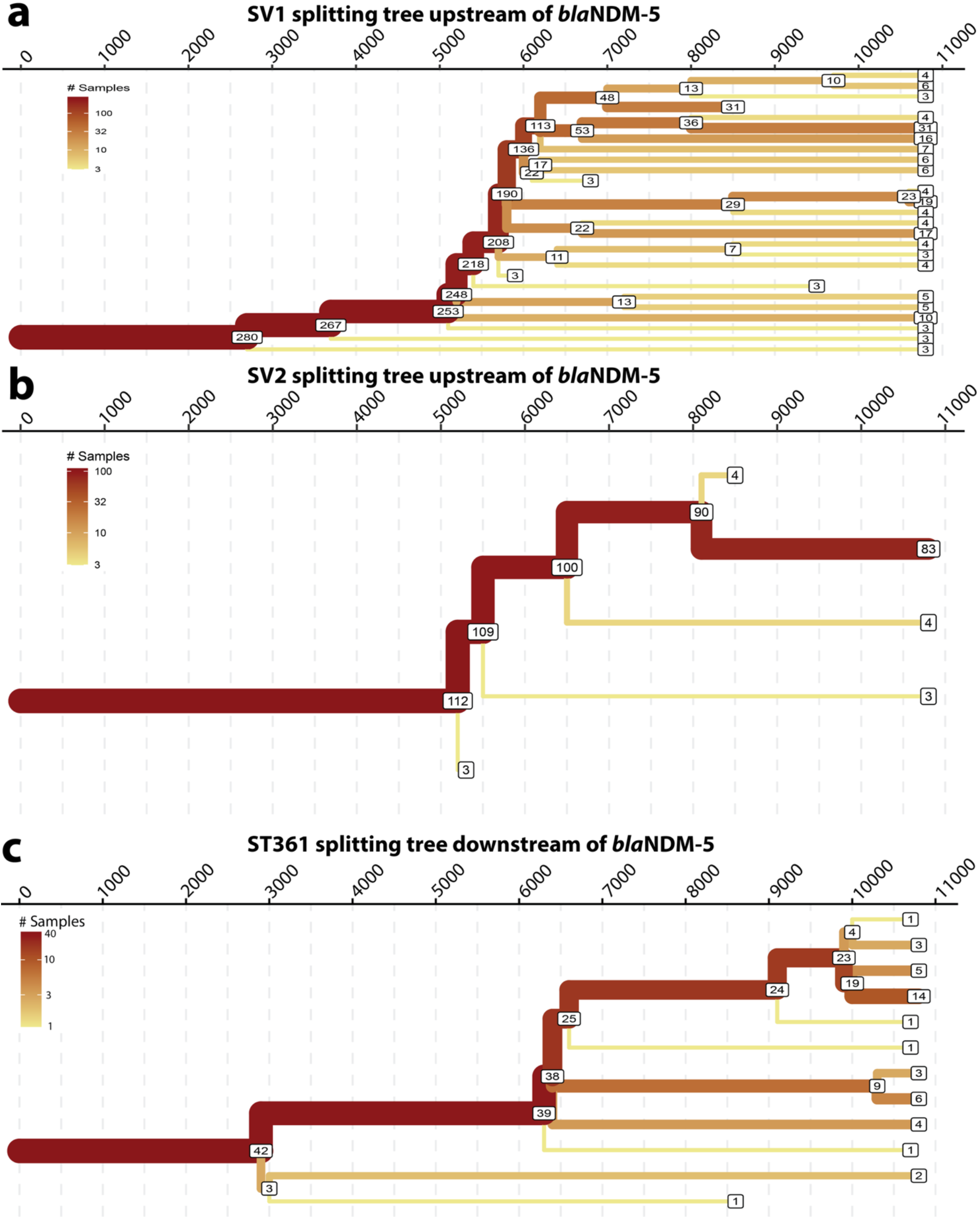
Splitting tree analysis upstream of *bla*_NDM-5_ and downstream in ST361. (**a,b**) A splitting tree analysis of the upstream region of *bla*_NDM-5_ for SV1 (a) and SV2 (b), the x-axis represents distance in base pairs from *bla*_NDM-5_. (**c**) A splitting tree of the structural variants identified downstream of *bla*_NDM-5_ in ST361 genomes. The labels on the node represent the remaining number of contigs on each branch. The x-axis represents distance from *bla*_NDM-5_.

**Supplementary Figure 9.**
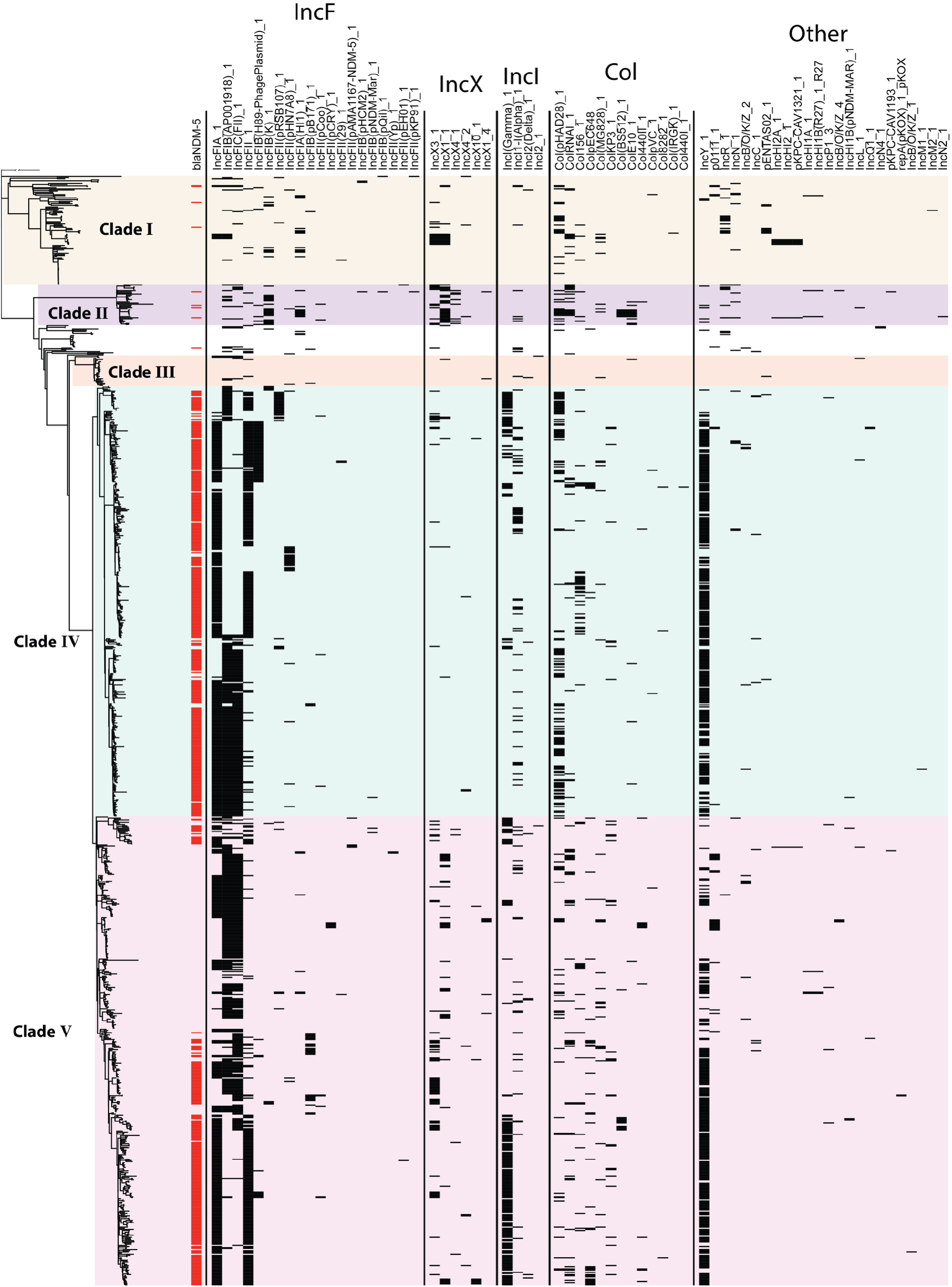
Heatmap of plasmid replicons on the ST361 phylogeny. Plasmid replicons were identified with the plasmid finder database using ABRicate. The red bars indicate genomes with *bla*_NDM-5_ present, while black bars indicate presence of plasmid replicons. Within each grouping of plasmid replicons, order is determined by prevalence.

The most common variant, which we designated structural variant 1 (SV1), accounted for 283/443 (63.9%) *bla*_NDM-5_ sequences. The SV1 downstream region comprised *bla*_NDM-5_, *ble*, *trpF*, *dsbD*, and ISCR2, followed by a partial class 1 integron containing *sul1* and *qacEdelta1* (Fig. 4a; Fig. 4b)^47^. In contigs containing SV1, two copies of IS*26* in the same orientation were frequently identified in the region flanking *bla*_NDM-5_ (Supplementary Fig. 7a). Although SV1 was predominantly associated with F plasmid replicons (241/262 contigs, with several contigs containing multiple copies of *bla*_NDM-5_), SV1 was also identified on contigs containing other plasmid replicons, including IncB/O/K/Z, IncI1, IncN, IncR, IncX, IncHI2, and IncY (Supplementary Data 8; Fig. 4c). The second structural variant (SV2), which accounted for 116/443 (26.2%) of *bla*_NDM-5_ downstream regions, comprised a conserved 4.7kb module containing *bla*_NDM-5_, *ble*, *trpF*, *dsbD*, IS*26, umuD* and part of IS*KOX3* (Fig. 4a; Fig. 4b). SV2 was predominantly associated with IncX3 plasmids (103/116 contigs; Fig. 4c; Supplementary Data 8), and its gene synteny was consistent with previous reports of *bla*_NDM_ variants on IncX3 plasmid backbones^48–50^. Notably, SV1 was less conserved than SV2, with IS elements occurring at variable positions relative to *bla*_NDM-5_ (Supplementary Fig. 7a). In contrast, the IS elements in SV2 were positioned at conserved sites, with 102/116 downstream regions maintaining nucleotide sequence homology and synteny extending >11kb downstream of *bla*_NDM-5_ (Fig. 4b; Supplementary Fig. 7b). The variable positioning of IS elements relative to *bla*_NDM-5_ in SV1 is consistent with greater structural diversity in the upstream region, where splitting-tree analysis revealed extensive fragmentation into numerous smaller branches (Supplementary Fig. 8a). In contrast, the upstream region of SV2 was largely intact, consistent with its conserved IS positioning and confinement to the IncX3 backbone (Supplementary Fig. 8b). Within ST361, SV1 was dominant, accounting for 39/42 (92.8%) sequences (Supplementary Fig 8c); of the remaining three sequences, two were identified on IncX3 plasmids and belonged to SV2, while the third was a unique structural variant on a contig without a predicted plasmid replicon.

To determine whether the plasmid backbone diversity supported independent conjugation acquisition events of *bla*_NDM-5_ containing plasmids in ST361, we extracted all *E. coli bla*_NDM-5_ positive plasmids from RefSeq (n = 402; see methods) and performed pling clustering. Pling groups plasmids into communities based on shared sequence content and further resolves these into subcommunities based on structural rearrangements^51^. This analysis identified one dominant plasmid community that encompassed 375/402 (93.3%) plasmids (Fig. 4c;

Supplementary Data 9). Within this community, multiple subcommunities were resolved, several of which contained ST361 plasmids alongside those from other high-risk lineages including ST167, ST410 and ST405 (Fig. 4c). Our analysis identified cointegrate and degenerate plasmids that bridged distinct plasmid backbones, consistent with on-going plasmid recombination and structural decay (Fig. 4c). Seven plasmid subcommunities, each containing distinct plasmid backbones, comprised more than 10 genomes distributed across multiple STs. Taken together, these findings suggest that the dissemination of *bla*_NDM-5_ in *E. coli* is driven by the conjugative transfer of several distinct F plasmids and a group of conserved IncX3 plasmids, and likely IS*26-*mediated mobilisation of a transposable unit carrying *bla*_NDM-5_ followed by clonal spread (Fig 4). Consistent with the plasmid backbone distribution observed in complete genomes, screening of plasmid replicons across the ST361 phylogeny identified diverse IncF replicon combinations associated with *bla*_NDM-5_ carriage, whereas IncX3 was associated with a small subset of *bla*_NDM-5_ positive genomes. These findings indicate that F plasmids are the predominant plasmid vehicles mediating *bla*_NDM-5_ dissemination within ST361 (Supplementary Fig. 9).

### *In silico* serotyping reveals a highly conserved *Klebsiella*-like serotype

To characterise the serotypes of ST361, we first analysed the colanic acid and O-antigen gene loci in complete ST361 genomes. This analysis revealed the absence of colanic acid biosynthesis genes, a defining feature of *E. coli* harbouring group 1 capsule^52^ (Fig. 5a). The genes encoding group 1 capsule are thought to have been acquired in *E. coli* from *Klebsiella* via recombination between the *galF* and *gnd* loci^53,54^. Accordingly, we used the *Klebsiella* primary capsule reference database implemented in Kaptive 3.0 to identify group 1 capsule loci in ST361^55^. This analysis identified 15 distinct group 1 capsule loci in the dataset, with the KL25^Group1^ (KL25^G1^) capsule locus predominant and present in 95.9% (941/981) of ST361 genomes (Fig. 2; Fig. 5b). Further analysis of O-and H-antigens with the *E. coli* EcOH database^56^ revealed most isolates carried the O9 O-antigen (902/981) and the H30 H-antigen (961/981). Interestingly, the O9 O-antigen shares high sequence similarity to the O3 O-antigen of *K. pneumoniae*^57^ (Fig. 5c). Collectively, these findings reveal a highly conserved KL25^G1^:O9:H30 serotype among ST361 isolates (882/981 genomes) that reflects cross-species convergence of a capsule and O-antigen polysaccharide serotype (Supplementary Data 2).

**Figure 5.**
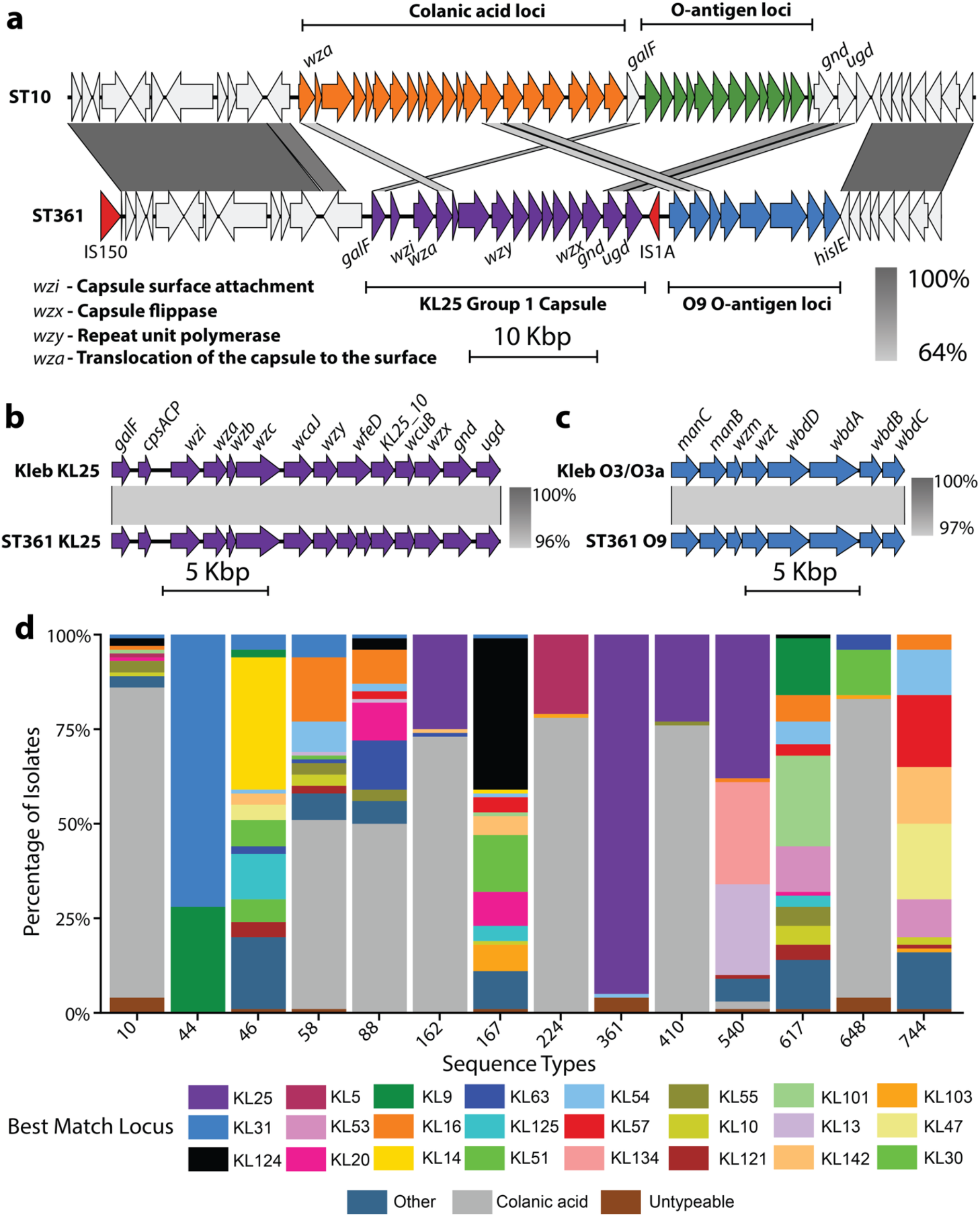
ST361 possesses a *Klebsiella*-like serotype architecture. (**a**) Pairwise DNA sequence alignment of a representative ST10 genome (NZ_CP060974.1) containing colanic acid and MS25536 (accession ID: JCANXH000000000) a representative ST361 genome with the KL25 group 1 capsule and O9 O-antigen loci. Genes coloured in orange represent the colanic acid locus, green indicates the O-antigen locus of the ST10 genome, purple defines the KL25 group 1 capsule locus, blue indicates the O9 O-antigen genes and red indicates insertion sequences determined by ISfinder. (**b**) Pairwise DNA sequence alignment of KL25 group 1 capsule from the Klebsiella reference database from Kaptive 3.0 and MS25536. (**c**) Pairwise DNA alignment of the *Klebsiella* O3/O3a locus from the *Klebsiella* O-antigen reference database and MS25536. (**d**) Group 1 capsule prevalence across *E. coli* sequence types. Sequence types with greater than 10% carriage of group 1 capsule are presented. Colours represent distinct group 1 capsule loci identified by Kaptive 3.0, with grey representing colanic acid.

While group 2 capsule diversity has been extensively studied in UPEC^58,59^, the extent of group 1 capsule diversity in *E. coli* remains largely undefined. We and others previously identified extensive group 1 capsule diversity within ST167^18,21^, prompting us to consider whether the conservation of a single group 1 capsule type is common in other *E. coli* STs or unique to ST361. To assess this, we screened 10,000 *E. coli* genomes from EnteroBase^20^, comprising 100 genomes from the 100 most common STs, collected between 2021 and 2026 with Kaptive 3.0^55^ (100STv2; Supplementary Data 10). This analysis identified 46 distinct group 1 capsule loci harboured in 921 genomes spanning 22 STs and revealed a broader distribution than previously reported, with most *E. coli* STs demonstrating greater group 1 capsule diversity than ST361 (Fig. 5d). The proportion of group 1 capsule containing genomes varied by ST, however, among the STs that harboured group 1 capsule there were several known CREC lineages, including ST10, ST167, ST617, ST744, ST410, ST648 and ST361 (Fig. 5d). Taken together, these results confirmed that the high conservation of KL25 across the ST361 lineage is characteristic of ST361.

### Virulence potential of ST361

To assess the virulence potential of ST361, we performed an *in silico* screen using a curated UPEC/ExPEC virulence factor database^18^. This analysis identified different virulence factors across clades (Supplementary Fig. 10a; Supplementary Data 11). Of note was the presence of iron acquisition systems *sitA* and *fecA* in clades IV and V that were largely absent in other clades (Supplementary Fig. 10a). However, ST361 harboured fewer virulence factors overall compared to lineages from phylogroups B2 and D, consistent with other phylogroup A and C lineages (Supplementary Fig. 10b). ST361 contained the type 1 fimbriae locus, a canonical UPEC virulence factor critical for bladder colonisation^60,61^. FimH typing revealed a conserved *fimH54* allele (Fig. Error! Reference source not found.). Analysis of the type 1 fimbriae locus in clades IV and V showed the presence of an IS*1A* element in clade IV genomes located between the end of the *fimE* coding sequence and the inverted repeat left (IRL) of the phase variable *fim* switch (*fimS*) region (Supplementary Fig. 11a; Supplementary Fig. 11b). To investigate if the IS element identified in clade IV impacted phase variation of the *fimS* region, we performed read mapping from short read sequencing of isolates from clades IV and V to the *fimS* region in the ON and OFF orientation using DISCus, a previously described method for evaluating *fim* switch orientation using genome sequence data^62^. This analysis showed evidence for phase variation of the type 1 fimbriae locus in strains from both clades IV and V, although the switch was largely in the OFF orientation (Supplementary Fig. 11c), suggesting the IS element present in clade IV genomes does not prevent inversion of the *fimS* region.

**Supplementary Figure 10.**
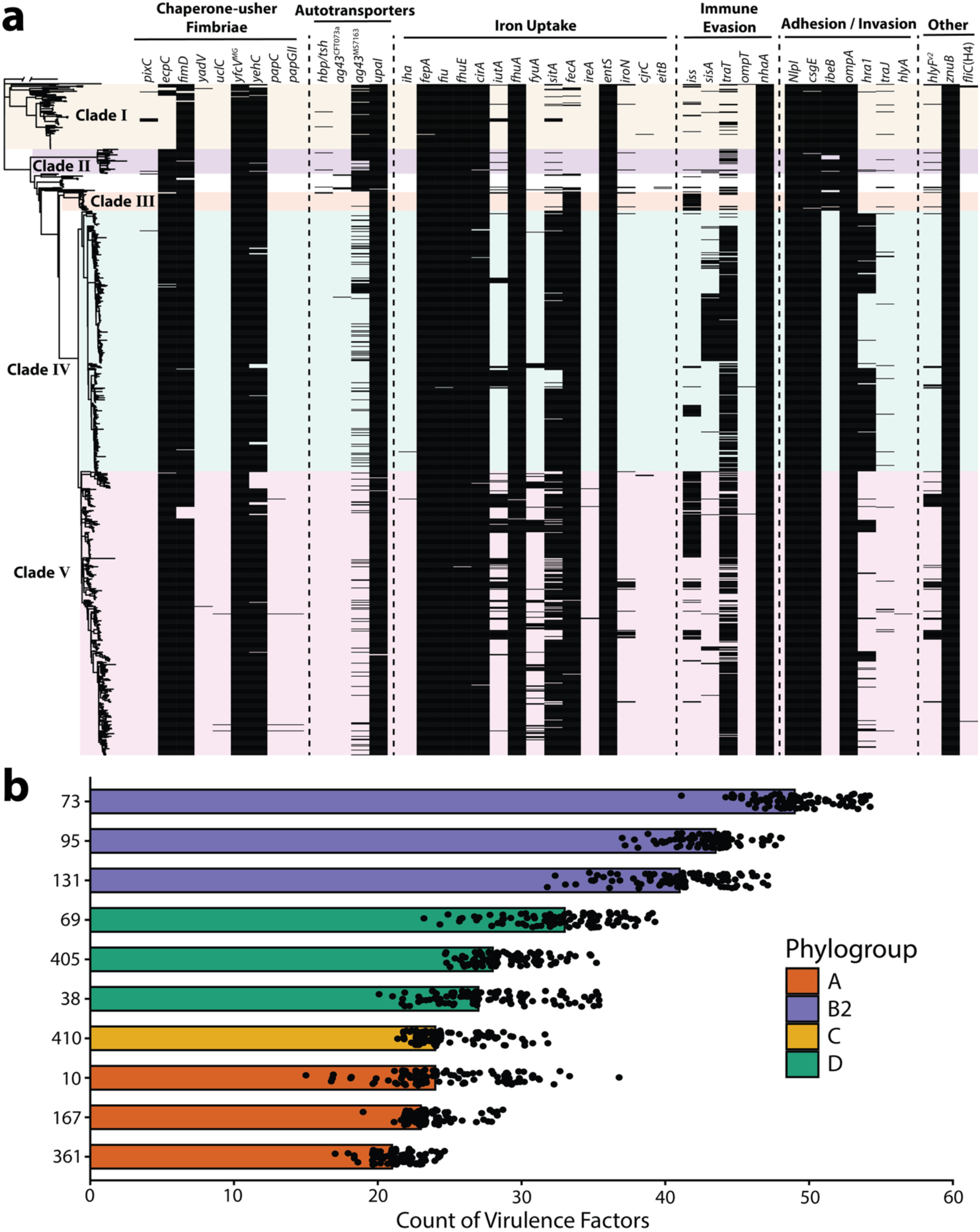
*In silico* virulence factor prediction of ST361 and other *E. coli* lineages. (**a**) Virulence factor heatmap of ST361 genomes. Presence of virulence factors is designated by black bars, whilst clades are designated by the coloured shading. (**b**) Median virulence factor score of selected *E. coli* lineages from phylogroups B2, D, C and A from the 100STv2. Dots depict the number of virulence factors identified in each genome using the UPEC/ExPEC virulence factor database with ABRicate. Colour of the bars represent the phylogroup.

**Supplementary Figure 11.**
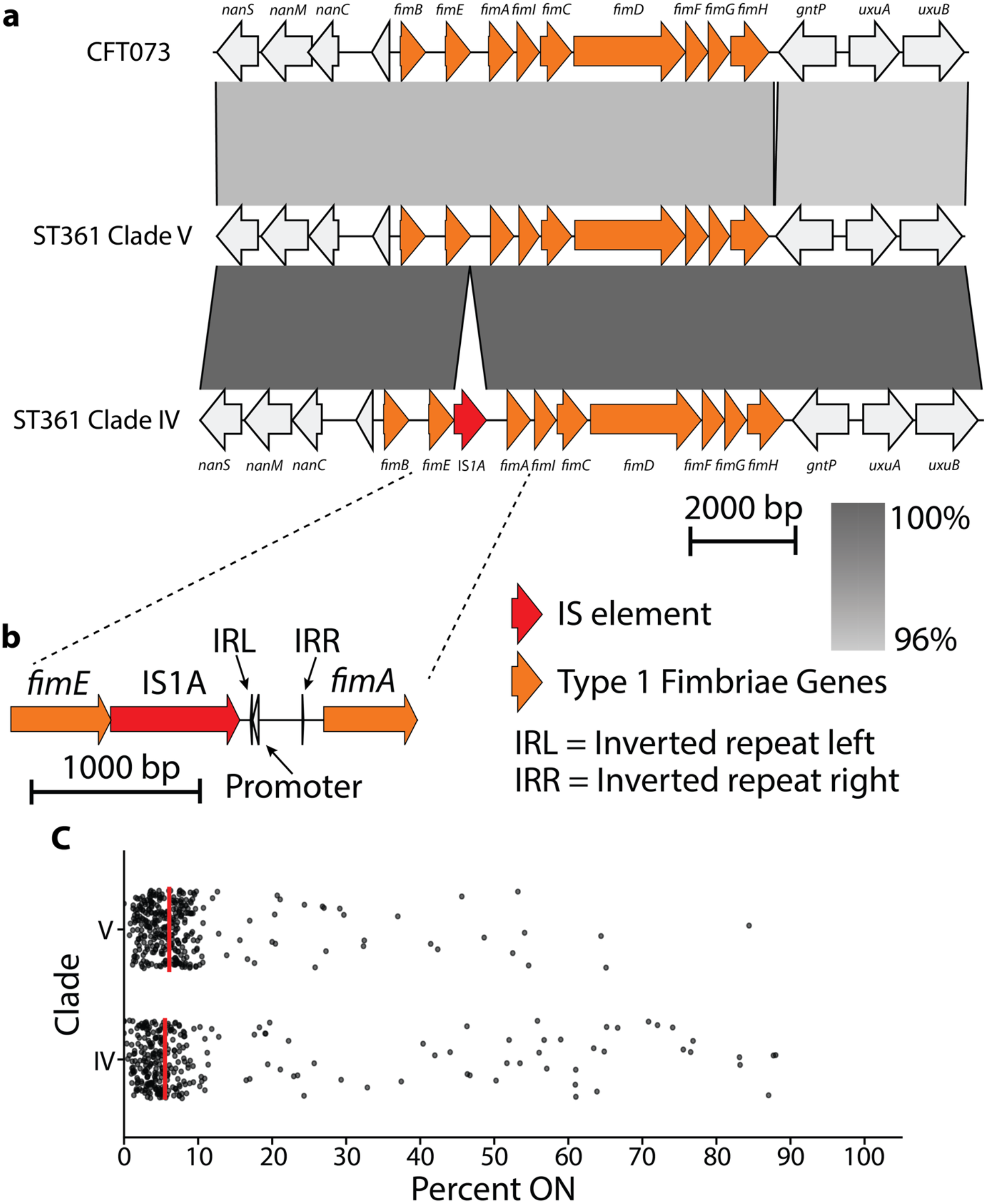
Analysis of type 1 fimbriae locus in ST361 clades IV and V. (**a**) Pairwise DNA sequence alignment of ST361 isolates from clades IV (MS25613) and V (MS25611) with CFT073. (**b**) Easyfig depiction of the *fimS* region and invertible promoter of type 1 fimbriae, an example of the phase OFF phenotype. (**c**) Read mapping of short read sequencing from 594 ST361 genomes to determine the proportion of reads that map to the phase ON orientation. Red bar indicates the median proportion of reads in the phase ON orientation.

### ST361 requires O-antigen for whole blood survival

Given the association of ST361 with UTI despite reduced UPEC-specific virulence factors, we assessed the virulence of ST361 isolates in a series of infection relevant *in vitro* and *in vivo* models. Representative strains of clade IV (MS25613) and clade V (MS25610) were selected based on gentamicin susceptibility to enable isogenic mutant generation (Supplementary Data 7), with capsule and O-antigen mutants generated using λ-Red mediated homologous recombination. To assess the contribution of the *Klebsiella*-like polysaccharide serotype to virulence of ST361 strains, we incubated wild-type, capsule-deficient and O-antigen-deficient mutants in fresh whole blood for one hour and assessed bacterial survival (Fig. 6a). Loss of the O-antigen significantly reduced survival of isolates from both clades (*P* < 0.01; Kruskal-Wallis test), whereas survival of the capsule-deficient mutants was comparable to that of their respective wild-type strains (Fig. 6b). Notably, wild-type ST361 strains exhibited survival in whole blood comparable to that of the prototypical UPEC phylogroup B2 strain CFT073 (ST73), which was included as a control.

**Figure 6.**
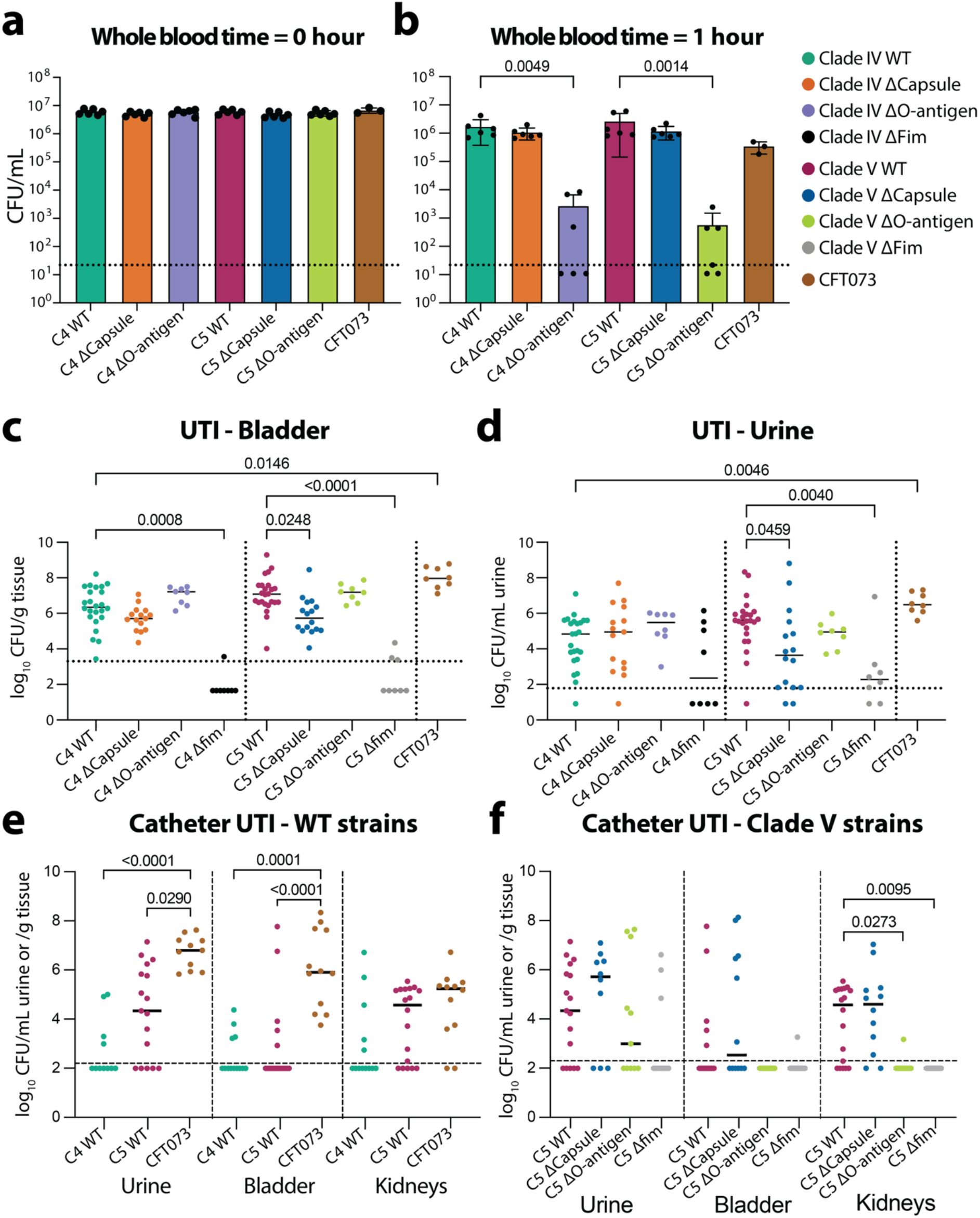
ST361 virulence assessment *in vitro* and *in vivo*. (**a,b**) Whole blood killing assay with bacterial CFU/ml plotted at both 0h (**a**) and 1h (**b**) incubation with human whole blood. Data is presented as mean ± (error) CFU in biological replicates (n = 6 human donors for ST361 wild-type and mutants, n = 3 human donors for CFT073). Statistical significance was determined by a Kruskal-Wallis test with Dunn’s multiple comparison post-hoc test. **(c,d)** Bacterial loads recovered from the (**c**) bladder and (**d**) urine of infected C57BL/6 mice expressed as median bacterial load per gram of tissue or mL of urine. Each symbol represents the bacterial load for an individual mouse at 24 h post infection. Representative clade IV (MS25613) and clade V (MS25610) strains and their isogenic mutants were used alongside UPEC control CFT073. Wild-type clade IV/V: n = 24 (3 pooled experiments); Δcapsule: MS25613 n = 15, MS25610 n = 16 (2 pooled experiments); ΔO-antigen: n = 8 (1 experiment); Δfim: n = 8 (1 experiment); CFT073: n = 8. Statistical significance was determined by Kruskal-Wallis test with Dunn’s multiple comparison post-hoc test. (**e,f**) Bacterial loads recovered from the urine, catheterized bladder, and kidneys of infected CBAJ mice expressed as median bacterial load per gram of tissue or mL of urine. Each symbol represents the bacterial load for an individual mouse at 24 h post infection. Representative clade IV (MS25613) and clade V (MS25610) strains and isogenic mutants of the clade V strain were used alongside UPEC control CFT073. CFT073: n = 12 (3 pooled experiments); wild-type clade IV: n = 13 (2 pooled experiments); wild-type clade V: n = 18 (3 pooled experiments); Δcapsule: n = 12 (2 pooled experiments); ΔO-antigen: n = 11 (2 experiments); Δfim: n = 11 (2 experiments). Statistical significance was determined by Kruskal-Wallis test with Dunn’s post hoc correction. All dotted lines represent the limit of detection. Only statistically significant comparisons are shown, with *P* < 0.05 considered statistically significant.

**Supplementary Figure 12.**
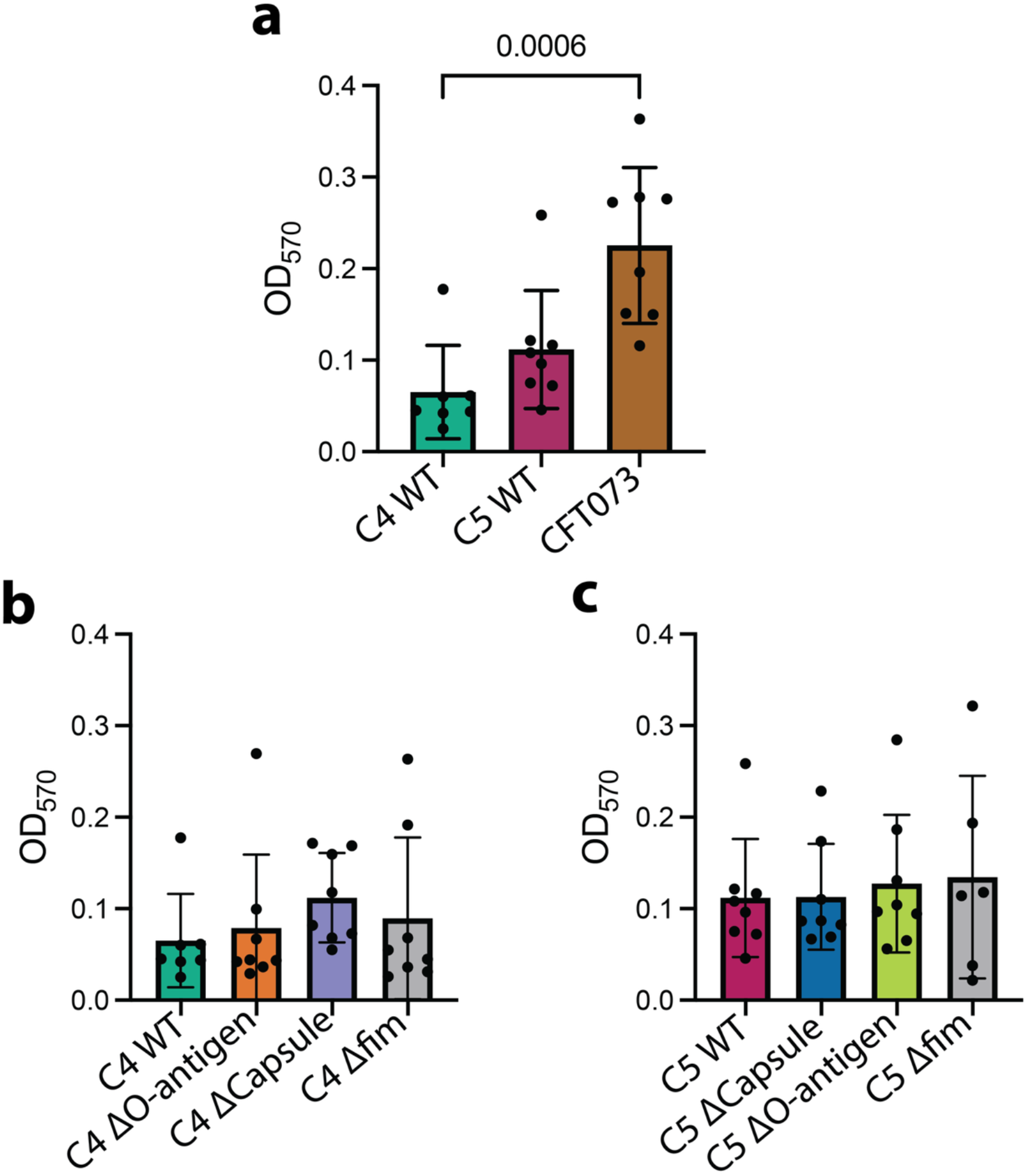
Biofilm formation on silicone Foley catheter segments. (**a-c**) Biofilm mass on silicone Foley catheter segments was measured by crystal violet staining after 24 h incubation in LB. Data is presented as mean crystal violet staining (measured at OD_570_), with each dot representing replicate catheter segments from 3-4 independent experiments. Statistical significance was determined by a Kruskal-Wallis test with Dunn’s multiple comparison post-hoc test. Only significant differences are noted.

### Representative ST361 clade IV and V isolates efficiently colonise the mouse bladder

Given the association of ST361 clades IV and V with UTI despite their reduced UPEC virulence factor repertoire, we sought to directly assess their uropathogenic potential using a murine UTI model. Wild-type strains MS25613 (clade IV) and MS25610 (clade V) efficiently colonised the bladder (Fig. 6c) and were recovered from urine at comparable levels (Fig. 6d). CFT073 colonised the bladder and urine at significantly higher levels compared to the clade IV strain (*P* = 0.0146) but not the clade V strain (*P* = 0.5213). We further demonstrated that mutation of the KL25^G1^ capsule resulted in an ∼1-log reduction in bladder colonisation for both strains which was only statistically significant in the clade V strain MS25610 (clade IV *P* = 0.5677; clade V *P* = 0.0248). In contrast, mutation of the genes encoding the O9 O-antigen did not significantly affect bladder colonisation (Fig. 6c). Mutation of the type 1 fimbriae genes resulted in attenuated colonisation of the bladder for both strains (*P* < 0.001; Fig. 6c). No statistically significant difference in the infection load of each mutant was observed in the urine for the clade IV strain, although the clade V strain showed significantly reduced colonisation of the urine in both the capsule (*P* = 0.0459) and type 1 fimbriae (*P* = 0.0040) mutants (Fig. 6d).

### The O-antigen and type 1 fimbriae are important determinants for kidney infection in a murine CAUTI model

Given the association between carbapenem resistance and healthcare settings, we compared the colonisation of the ST361 strains to CFT073 in a CAUTI model. In contrast to the UTI model, both wild-type strains MS25613 (clade IV) and MS25610 (clade V) colonised the urine and bladder poorly in this model (Fig. 6e). Bacterial burden in the kidneys was comparable across strains, although significantly fewer mice infected with the clade IV strain developed pyelonephritis compared to those infected with CFT073 (38.5% v 83.3%, *P* = 0.0414 by Fisher’s Exact test; Supplementary Data 12). Only the clade V strain achieved sufficiently robust colonisation in this model to allow the contributions of the conserved KL25^G1^ capsule, O9 O-antigen and type 1 fimbriae to be examined. Mutation of the KL25^G1^ capsule had no impact on bacterial burden or incidence of organ colonisation in the CAUTI model (Fig. 6f). Mutation of O-antigen significantly reduced bacterial burden in the kidneys (Kruskal-Wallis *P* = 0.0273) and significantly decreased the overall incidence of kidney colonisation (Fisher’s Exact *P* = 0.0008; Supplementary Data 12). Similarly, mutation of type 1 fimbriae significantly reduced bacterial burden in the kidney (Kruskal-Wallis *P* = 0.0095) and the incidence of kidney colonisation (Fisher’s Exact *P* = 0.0001).

Since biofilm formation on the catheter is a hallmark of CAUTI, we further assessed biofilm formation on Foley catheter segments (Supplementary Fig. 12). The ST361 strains both exhibited reduced catheter biofilm formation compared to CFT073, which was the most pronounced for the clade IV strain (*P* = 0.0006) compared to the clade V strain (not significant; *P* = 0.065). However, loss of either capsule, O antigen, or type 1 fimbriae had no impact on biofilm biomass for either strain. Decreased catheter biofilm formation could therefore contribute to the decreased burden of these strains in the urine and catheterized bladder but likely does not contribute to the further defects observed for the clade V mutants in the murine CAUTI model.

## Discussion

This study identified ST361 as an emerging high-risk lineage of CREC associated with UTI in line with previous reports^32,34,35,43^. Phylogenomic characterisation of ST361 identified two dominant clades marked by extensive carbapenem resistance and a highly conserved *Klebsiella*-like capsule and O-antigen serotype that contribute to pathogenesis. Bayesian phylodynamic analyses support the recent emergence and intercontinental spread of these clades, with the dissemination of the primary carbapenem resistance gene *bla*_NDM-5_ likely driven by both IS*26*-mediated transposition and plasmid conjugation.

ST361 clade IV and clade V exhibit a near pan-drug resistance profile, with carbapenem resistance driven primarily by *bla*_NDM-5_. The co-occurrence of *bla*_NDM-5_ with the YRIN insertion in the PBP3 binding protein (FtsI_N337NYRIN) also compromises recently developed antibiotics such as cefiderocol and aztreonam-avibactam that are used to treat carbapenem-resistant *Enterobacterales*^6,7,63^. The clinical relevance of this resistance profile is reflected in the consistent representation of ST361 within the Centers for Disease Control and Prevention (CDC) National Healthcare Safety Network (NHSN; BioProject PRJNA288601), a nationwide surveillance program that sequences carbapenem-resistant bacteria from healthcare-associated infections (HAI)^64–66^. Within this HAI dataset, 45.1% (170/377) of ST361 isolates were derived from urinary sources, consistent with sustained transmission of this multidrug-resistant lineage in healthcare environments where carbapenem exposure is highest.

Bayesian phylodynamic analysis dated the emergence of ST361 clade IV and clade V to approximately 2006, with Asia as the inferred geographical origin within the sampled dataset. This temporal emergence coincides with increased global consumption of carbapenem antibiotics; between 2000-2010, carbapenem usage increased approximately 45% across 71 countries globally^67^. Increased carbapenem use, particularly in hospital settings, may provide selective pressure favouring the emergence and global dissemination of *bla*_NDM-5_ harbouring ST361 isolates. The phylogeographic analyses support multiple independent transition events of ST361 between continents, providing evidence for ongoing intercontinental dissemination.

Previous studies have shown that *bla*_NDM-5_ and other *bla*_NDM_ alleles can be mobilised by IS*26* elements and inserted into diverse plasmid backbones^46,68–72^. By integrating these observations with our large-scale analysis of ST361 and RefSeq plasmids^73^, we build on existing knowledge to further understand the molecular mechanisms underlying the spread of *bla*_NDM-5_. We identified two conserved structures (SV1 and SV2) downstream of *bla*_NDM-5_ and found that *bla*_NDM-5_ was carried on predominantly IncX3 and F plasmid backbones across high-risk CREC lineages, including ST361, ST167, ST410 and ST405. These findings indicate that dissemination of *bla*_NDM-5_ across lineages is associated with the spread of conserved *bla*_NDM-5_-associated structures on a limited range of plasmid backbones. The evidence suggests that SV1 may be mobilised as a translocatable unit by the flanking IS*26* elements, consistent with its organisation as a pseudo-compound transposon^74–76^, although this remains to be experimentally demonstrated. Indeed, the reduced conservation of SV1, along with its presence on plasmids with diverse replicons/backbones, is consistent with ongoing transposition and/or structural rearrangement of the *bla*_NDM-5_ region. In contrast, the strong conservation of SV2 and its restriction to IncX3 plasmids suggests that the dissemination of SV2 is primarily driven by IncX3 plasmid spread rather than frequent transposition. This multi-layered model of IS-mediated acquisition, clonal expansion and plasmid conjugation provides a framework to understand the rapid dissemination of carbapenem resistance in *E. coli*.

ST361 is distinguished by genes encoding a highly conserved KL25^G1^:O9:H30 serotype, sharing the KL25^G1^ capsule type with *K. pneumoniae*, while the O9 O-antigen is antigenically identical to the O3 O-antigen locus in *K. pneumoniae*^57,77^. The high similarity of both loci to their *K. pneumoniae* counterparts demonstrates convergence of serotype-defining surface polysaccharides in different pathogens. Within *E. coli* population studies and vaccine design, group 1 capsule is relatively understudied. However, recent developments in capsule-targeted interventions in *K. pneumoniae*, including monoclonal antibodies and vaccines^78,79^, raise the possibility of cross-protection against CREC lineages encoding related capsule types. Such strategies would also help to address the problem of diminishing effective antibiotics for near-pan-resistant isolates in ST361 and other CREC clones.

ST361 harbours reduced UPEC specific virulence factors compared to classical UPEC lineages such as ST73, ST95 and ST131, consistent with its phylogroup A background. Despite this reduction, isolates from clades IV and V colonised the bladder and established infection in a murine model of UTI, albeit at lower levels than the prototypical UPEC strain CFT073. We propose that ST361 represents an atypical uropathogen whose success in healthcare settings is driven primarily by antimicrobial resistance rather than enhanced virulence. In this model, the near-pan drug resistance of ST361 imparts a fitness advantage in hospital settings where antibiotic pressure is highest, enhancing ST361 persistence and transmission where classical (and more susceptible) UPEC lineages are eliminated^80,81^. Our experiments revealed the *Klebsiella*-like capsule and O-antigen both contributed to the pathogenesis of ST361 in niche-specific contexts. The KL25^G1^ capsule contributed to ST361 bladder colonisation, while the O9 O-antigen was important for survival in human whole blood and colonisation of the kidneys in a CAUTI model. The differences observed between the UTI and CAUTI models employed in this study may reflect differences in innate immune responses between the two models. Beyond differences in the type of infection, the infection models also used different mouse strains; C57BL/6 mice for the UTI experiments and CBA/J mice for the CAUTI experiments. These mice were used to reflect our established models^82,83^ and enabled direct comparison of ST361 phenotypes to the UPEC reference strain CFT073. Future work will evaluate the pathogenicity of our ST361 strains using the same mouse strain across both infection models. In addition, since many CAUTIs are polymicrobial^84–86^, it would be of interest to investigate whether ST361 catheter infection is enhanced in the presence of other pathogens.

This study has several limitations. (i) The geographic distribution of ST361 isolates in EnteroBase^20^ is heavily skewed towards North America, with 86.2% of isolates deposited between 2021-2025 originating from this region. The majority of these isolates were contributed by the CDC NHSN surveillance programme. This reflects a broader characteristic of EnteroBase^20^, where depositions inevitably mirror the sequencing priorities of contributing organisations rather than a structured global surveillance dataset. Consequently, the frequency of ST361 depositions should not be interpreted as a direct measure of prevalence or clinical incidence. Instead, ST361 should be considered in comparison to other CREC lineages rather than UPEC more broadly. Consistent with this, the clinical significance of ST361 extends beyond North America, with multiple national and single-centre surveillance studies across diverse geographic regions independently identifying ST361 as a large contributor to the global CREC burden (Supplementary Data 1)^16,21,32–39^. (ii) The phylogeographic analyses are constrained by the same geographic biases. As such, the inferred geographical origin of Asia should be interpreted within the context of the analysis that restricted the model to isolates from North America, Europe and Asia due to sparse sampling in other regions of the world. (iii) The proposed model of *bla*_NDM-5_ dissemination is based upon genomic inference rather than experimental validation. Although consistent with previous studies describing IS*26*-mediated mobilisation, the dynamics of *bla*_NDM-5_ acquisition and plasmid conjugation were not directly assessed in this study. (iv) The *in silico* serotyping methods used in this study demonstrate high DNA sequence similarity between the capsule and O-antigen in *K. pneumoniae* and point towards cross-species convergence of surface polysaccharides. However, cross-species reactivity of antibodies for the *Klebsiella* capsule or O-antigen were not tested.

In conclusion, our data support ST361 as an emerging CREC clone associated with UTI, driven by clades IV and V. The acquisition of a *Klebsiella-*like capsule and O-antigen serotype and plasmids that confer carbapenem resistance, are likely key drivers in this atypical phylogroup A *E. coli* lineage. The near-pan drug resistance of ST361, coupled with its intercontinental spread, highlights the need for enhanced global surveillance of this emerging pathogen.

## Methods

### Ethics approval

Mouse UTI experiments were approved by the University of Queensland Animal Ethics Committee (approval number 2022/AE000469). Mouse catheter-associated UTI experiments were approved by the Institutional Animal Care and Use Committee of the University at Buffalo (approval number MIC31107Y), in accordance with the Office of Laboratory Animal Welfare, the U.S. Department of Agriculture, and the Association for Assessment and Accreditation of Laboratory Animal Care. Mice were anesthetized with a weight-appropriate dose (0.1 ml for a mouse weighing 20 g) of ketamine/xylazine (80 to 120 mg/kg ketamine and 5 to 10 mg/kg xylazine) by intraperitoneal injection and euthanized by CO2 with vital organ removal. Fresh peripheral blood was collected from healthy human donors following informed consent and with approval from the University of Queensland Human Ethics Committee (approval numbers 2025/HE001619).

### EnteroBase metadata

The EnteroBase^20^ *E. coli* dataset was accessed on the 17^th^ of December 2025. To identify CREC lineages from urinary sources we first filtered the dataset for isolates from human sources (search terms: ‘Human|Homo sapiens’) followed by urinary sources (search terms: ‘urine|UTI|urinary’), yielding 20,331 genomes. Draft genome assemblies for these isolates were downloaded directly from EnteroBase^20^ and screened for antimicrobial resistance with AMRfinderplus^87^ v3.11.20. To determine the proportion of CREC isolates from non-urinary human sources, we applied the same filters as before but excluded isolates from urinary sources, yielding 127,564 isolates. Given the size of this dataset, we retrieved pre-computed Amrfinderplus annotations available within EnteroBase^20^ rather than screening assemblies independently.

### Phylogenetic analysis

Publicly available ST361 genomes were downloaded directly from EnteroBase^20^. Quality control with QUAST v5^88^ was performed on draft assemblies, assemblies that contained more than 500 contigs and 20 undetermined bases per 100kb were removed. In total, 981 genomes were curated to produce the maximum likelihood phylogenetic tree. Briefly, we employed snippy and snippy-core v4.6.0 (https://github.com/tseemann/snippy) to call core-genome SNPs and generate a pseudo-genome alignment with SNPs reinserted into reference strain MS25613 (accession ID: JCANXK000000000). Gubbins v3.2.1^89^ with default parameters was used to filter recombination regions. IQ-TREE v2.1.2^90^ utilising the GTR+G model and 1,000 ultrafast bootstraps^91^ was used to generate the maximum likelihood phylogenetic tree. ED1a (accession ID: CU928162) was used as an outgroup for phylogenetic rooting. To determine the phylogenetic clade structure of ST361 we implemented Fastbaps v1.0.8^40^. The phylogenetic tree was visualised in R using ggtree^92^.

### Accessory genome analysis

The *in silico* AMR profile of ST361 genomes was determined using AMRfinderplus^87^. Group 1 capsule loci (and colanic acid which was manually added) were identified and typed using Kaptive v3.0^55^ with the *Klebsiella* primary reference database^93^; *E. coli* O-antigen was determined using ABRicate v0.8 with the EcOH database^56^. DNA pairwise synteny of the capsule and O-antigen loci were determined using EasyFig^94^. The sequence type of complete genomes was determined using the mlst software v2.11 (https://github.com/tseemann/mlst). ISfinder^95^ was used to identify IS elements presented in EasyFig figures. Clermontyper was used to determine phylogroups^96^. Fimtyper was used to determine the *fimH* allele of ST361 genomes^97^.

### 100STv2 database

The EnteroBase^20^ *E. coli* dataset was filtered for isolates collected between 01/01/2021 and 12/11/2025. A random selection of 100 genomes from each of the 100 most common sequence types deposited to EnteroBase^20^ during this period. This resulted in a collection of 10,000 genomes that we designated as 100STv2.

### Virulence factor analysis

Carriage of UPEC-specific virulence factors was determined by screening with the UPEC specific virulence factor database^18^ using ABRicate v0.8. To compare virulence factor carriage across both lineages and phylogroups we screened selected *E. coli* STs for their carriage of UPEC-specific virulence factors. Representative isolates (n = 100) of ST361, ST10, ST167, ST69, ST405, ST38, ST73, ST95 and ST131 were retrieved from the 100STv2 database.

To determine the proportion of isolates from clades IV and V in the phase ON or OFF orientation for type 1 fimbriae expression we used DISCus, a previously described method^62^. Briefly, publicly available short-read sequencing data for each isolate were mapped to a pseudo-reference sequence containing the *fimS* invertible switch in both orientations with 1kb flanking regions included. Read counts that mapped to the borders of each orientation were obtained (A_1 and A_2 for the OFF orientation; B_1 and B_2 for the ON orientation). To ensure accurate quantification of read mapping, isolates were excluded from the analysis if neither orientation was supported by a minimum of 20 reads at both border regions (e.g. A_1 < 20 and A_2 < 20 or B_1 < 20 and B_2 < 20). The percentage of reads in the ON orientation were calculated and plotted.

### Generation of isogenic mutant strains

To generate isogenic mutants of the capsule, O-antigen and type 1 fimbriae in ST361 strains MS25610 (clade V) and MS25613 (clade IV) we used a λ-Red recombinase system^98^. To mutate the capsule in ST361 we designed a construct that replaced genes between *cpsACP* and *gndA* with the chloramphenicol resistance gene. The construct consisted of ∼500 bp PCR products of the upstream and downstream region of the capsule loci, fused together with the chloramphenicol resistance gene isolated from the pKD3 vector in a three-way PCR. To mutate the O-antigen and type 1 fimbriae in ST361, we targeted *wbdC* (O-antigen) and *fimD* (type 1 fimbriae). To insertionally inactivate *wbdC* and *fimD* with the chloramphenicol cassette we amplified the chloramphenicol cassette with primers that contained 50 bp overhangs of the immediate upstream and downstream region of the specified genes. All constructs were electroporated into electrocompetent λ-Red expressing cells, where homologous recombination replaced the target sequences. All mutant strains were confirmed by sequencing.

### Antibiotic susceptibility testing

Antibiotic susceptibility testing of ST361 isolates was performed using broth microdilution (GN7F plates Sensititre; Thermo Fisher Scientific) and results were interpreted according to European Committee on Antimicrobial Susceptibility Testing (EUCAST) clinical breakpoints criteria V16.0^99^. For antibiotics that do not have a defined breakpoint according to EUCAST guidelines (e.g. minocycline) we deferred to Clinical and Laboratory Standards Institute (CLSI) guidelines^100^. For cefiderocol MIC testing by broth microdilution, iron-depleted cation-adjusted Mueller-Hinton broth (ID-CAMHB; In-house preparation) was used according to CLSI and manufacturer recommendations. Isolates were subcultured from storage on to 5% horse blood agar for 18-24 h at 37 °C prior to preparation of a 0.5 McFarland solution in ID-CAMHB for a final inoculum of 5 x 10^5^. Plates were inoculated once for each isolate and incubated at 37 °C for 18 to 20 h prior to reading. One positive growth control well and one negative control well were included in each plate. Quality control of cefiderocol was performed using *E. coli* ATCC 25922 and *P. aeruginosa* ATCC 27853.

### Whole blood killing assay

Fresh blood from healthy donors was collected using Vacuette tubes containing sodium heparin (Greiner Bio-one). Blood (90µl) was inoculated with 10µl of bacteria (∼10^7^ CFU) in a 96-well plate and incubated at 37°C for 1 h. Bacteria were enumerated to determine CFU prior to incubation with whole blood (0 h) and at 1 h post-incubation.

### Mouse infection experiments

All experiments were performed using the following housing conditions: (light:dark cycle 12:12 h, room temperature 21 ± 1 °C, humidity 50 ± 10%). The mouse UTI model was performed essentially as previously described^83,101^. All strains were enriched for type 1 fimbriae expression by performing three consecutive 24 h static growth cycles in low-salt LB (5 g/L). Strains were cultured overnight in LB broth statically for inoculum preparation. Female C57BL/6 mice (8-10 weeks) were infected via transurethral inoculation at ∼5×10^8^ CFU per 20-30 µL. Mice were euthanised by cervical dislocation at 24 h post infection. Bacterial loads corresponding to each strain in the urine, bladder and kidney of infected mice were enumerated by plating onto LB agar.

For the mouse CAUTI model, strains were enriched for type 1 fimbriae expression as above and female CBA/J mice (Jackson Laboratory, 6-8 weeks) were anesthetized via IP injection with ketamine/xylazine (80-120 mg/kg ketamine and 5-10 mg/kg xylazine) and transurethrally inoculated with ∼1×10^5^ CFU per 50 µL. A 4 mm segment of sterile silicone tubing (0.64 mm O.D., 0.30 mm I.D., Braintree Scientific Inc.) was advanced into the bladder during inoculation to be retained for the duration of the study as previously described^82^. After 24 h, mice were euthanized by CO_2_ with vital organ removal and bacterial loads corresponding to each strain in the urine, catheterized ladder, and kidney of infected mice were enumerated as above.

### Catheter biofilm experiments

Size 14 French silicone Foley catheters were aseptically sectioned into 15 mm segments, and one segment was wedged into each well of 24 well plate to remain fully submerged in 1 ml of LB. Each well was inoculated with ∼1×10^7^ CFU of the indicated strain and incubated for 20 h. Catheter segments were gently washed in PBS, air-dried, stained with 0.1% crystal violet, and washed again to remove excess stain. The crystal violet was then solubilized by vortexing the catheter segment in 95% ethanol and absorbance (OD_570_) was measured in a BioTek Synergy H1 plate reader^102^.

### Analysis of *bla*_NDM-5_ and plasmid diversity

To investigate the structural conservation of the *bla*_NDM-5_ gene across *E. coli* we compiled a dataset of 5,052 complete genomes from RefSeq (accessed 22^nd^ of May 2026; that included 55 complete ST361 genomes). The *bla*_NDM-5_ gene was identified in each genome by a BLASTn^103^ search, and 10 kb downstream flanking sequence was extracted. The extracted downstream sequences were compared using a splitting tree approach^46^, which partitions contigs based on shared homology at increasing distances from *bla*_NDM-5_. Splits into structural variants were defined when both daughter branches contained at least 10% of the parent population. We further extracted 10 kb of the upstream flanking sequence of *bla*_NDM-5_ in genomes belonging to the two structural variants. IS elements in the flanking regions relative to *bla*_NDM-5_ were identified using ISEScan v1.7.3^104^.

To characterise the plasmid diversity among *bla*_NDM-5_ carrying isolates, we extracted all contigs from long-read assemblies and retained those between 10 kb and 500 kb to exclude fragmented sequences and chromosomal contigs. Contigs identified to contain a plasmid replicon by the PlasmidFinder database within ABRicate v0.8 were retained for downstream analyses. We used pling^51^ with a containment distance of 0.3 and a DCJ distance ≤ 4 to identify more recent transmissions. A network was constructed in R using igraph, in which nodes represent plasmids and edges connect pairs within a community that had a DCJ distance ≤ 4. The network was visualised using ggraph with a Fruchterman-Reingold layout. All plasmid sequences were annotated with Bakta v1.5.1^105^.

### Nanopore sequencing

To produce complete genome sequences, we performed long-read sequencing in two ways, (1) in-house and (2) by outsourcing nanopore sequencing to plasmidsaurus (https://plasmidsaurus.com/technical-documentation/genome). The method of sequencing and sequencing metrics are listed in Supplementary data 13. For our in-house sequencing we used an Oxford Nanopore R10.4.1 flow cell with the Rapid Barcoding Kit 24 V14 (SQK-RBK114.24) and bases called with high accuracy base calling. Reads were filtered using Filtlong v0.2.1 to remove the bottom 5% of fastq reads and assembled using Autocycler v0.5.2^106^ with all available assemblers. All assemblies were subsequently polished with Polypolish^107^ v0.6.0 using available Illuminia short-read data.

### Phylodynamic analysis

For the Bayesian phylodynamic analysis of clades IV and V, we curated a dataset of 700 genomes comprised of 350 genomes from each clade. This dataset was randomly sampled from the full dataset with selection proportional to the yearly prevalence of each clade to retain the temporal structure of the dataset. Generation of a recombination-filtered core-genome SNP alignment (see Phylogenetic analysis in Methods) were generated using MS25613 (accession ID: JCANXK000000000) as the reference genome. Tip dates of the genomes were defined by collection year and month when available; genomes lacking a collection month were assigned to January of the collection year, which remains robust for phylodynamic analysis of bacteria over longer sampling intervals^108^. To investigate if the data contained a temporal signal we performed a Bayesian evaluation of temporal signal (BETS)^109^ using BEAST (v1.10.4). Briefly, this method compares the fit of heterochronous data (real tip dates) and isochronous data (all tips with the same date) to assess if the dataset contains a temporal signal. We fit a coalescent model with constant effective population size and a strict molecular clock with an MCMC chain of 1 x 10^9^ iterations. We used generalised stepping stone sampling with 100 steps and a 1 x 10^8^ MCMC steps to estimate the marginal likelihood, which is used to compare the model with heterochronous and isochronous tip dates. The BETS analysis supported the existence of temporal signal with a decisive log Bayes Factor of +738 for the heterochronous tips.

Bayesian phylogenetic analyses were subsequently performed using BEAST v1.10.4 with a monophyletic constraint applied to clade V. We used a GTR substitution model with empirical base frequencies and gamma-distributed rate heterogeneity with four rate categories (GTR+G4). An uncorrelated relaxed clock^110^ was selected over a strict clock based on the posterior coefficient of variation, which did not abut zero (Mean 0.911 [95% HPD 0.7979, 1.0307]). The relaxed clock rate was assigned a gamma prior (shape = 2, scale = 3.0 x 10^-7^, initial value = 1.0 x 10^-5^), informed by previous studies^28,31,111^. A time-aware GMRF Bayesian Skyride coalescent model was used as the tree prior^42,112^. The MCMC chain was run for 2 x 10^9^ iterations, with the first 10% discarded as burn-in. Convergence was assessed in both Tracer v1.7.2 and Beastiary^113^ with all parameters achieving an effective sample size (ESS) > 200. A maximum clade credibility tree was generated using TreeAnnotator with 10% burn-in applied. Individual Bayesian phylogenetic analyses were performed separately for clades IV and V using the same model settings and convergence criteria. For the clade V analysis MS25610 (accession ID: JCANXN000000000) was used as reference, while MS25613 was used as reference for clade IV. MCMC chains were run for 1 x 10^9^ iterations, sampling every 100,000 samples.

To reconstruct the geographical dynamics of clades IV and V, we performed a discrete phylogeographic analyses separately for each clade. We used the continent of collection as the discrete trait and included North America, Europe and Asia as a roughly-equal amount of genomes originated from these regions, reducing potential sampling bias^114^. Due to the high proportion of North American genomes between 2021 to 2025, we restricted the analysis to isolates collected before 2022. This resulted in a more geographically balanced dataset spanning seven years for clade IV (n = 119 genomes) and fourteen years for clade V (n = 155 genomes). Analyses were performed under the same model as above. A strict clock was preferred over the relaxed clock likely due to the reduced dataset. The same clock prior was applied as described previously. An asymmetric discrete trait substitution model with Bayesian Stochastic Search Variable Selection (BSSVS) was applied to the location (the discrete trait). Markov jump counts were used to estimate the number of transition events between continents. MCMC chains were ran for 1 x 10^9^ iterations, sampling every 100,000 iterations and the first 10% of the chain discarded as burn-in. ESS of all parameters were > 200 to ensure convergence. The mean number of transition events across the MCMC chain were displayed as a chord diagram.

### Statistical analyses

To determine statistical significance for *in vitro* and *in vivo* data we used GraphPad Prism (version 10.2.2). All other statistical analyses were performed in R. Information regarding statistical tests used and sample sizes are referenced in the appropriate text and/or figure legends. No statistical method was used to determine sample size. No data was excluded from the analyses. The investigators were not blinded to allocation during experiments and assessment.

## Acknowledgements

This work was supported by an Australian National Health and Medical Research Council (NHMRC) Ideas grant to MAS, M-DP and NTKN (2048896) and an NHMRC Ideas grant to MAS and M-DP (2037698). BMF is supported by a University of Queensland Health Research Accelerator grant. LAF is supported by the Medical Research Future Fund (GA167770). We thank the Centers for Disease Control and Prevention (CDC) National Healthcare Safety Network (NHSN), as well as other surveillance networks, for making genome sequence data used in this study publicly available. This research was also supported by use of the Nectar Research Cloud and by QCIF. The Nectar Research Cloud is a collaborative Australian research platform supported by the [NCRIS](https://www.education.gov.au/ncris)-funded [Australian Research Data Commons](https://ardc.edu.au/?nowprocket=1) (ARDC).

